# SMFNet reveals principles of combinatorial transcription factor assembly

**DOI:** 10.64898/2026.10.30.762954

**Authors:** Till Schwämmle, Nathaniel Diamant, Yuliya Badayeva, Rozemarijn Kleinendorst, Anshul Kundaje, Arnaud R. Krebs

**Affiliations:** Genome Biology Unit, EMBL Heidelberg, Meyerhofstraße 1, 69117 Heidelberg, Germany; Department of Computer Science, School of Engineering, Stanford University; Department of Genetics, School of Medicine, Stanford University; Faculty of Biosciences, Collaboration for Joint PhD Degree between EMBL and Heidelberg University, 69120 Heidelberg, Germany

## Abstract

Combinatorial assembly of transcription factors (TFs) opens chromatin at cis-regulatory elements (cREs) to establish cell-type-specific gene expression. Dissecting the role of individual TFs and their collective function has been limited by the diversity of TF binding-site arrangements across cREs. A single regulatory sequence can support multiple configurations of TF and nucleosome occupancy, raising the possibility that distinct chromatin states reflect different TF assemblies. Predictive models linking regulatory sequence to chromatin accessibility collapse these distinct molecular states into population averages, obscuring how TF assemblies shape accessible chromatin. Here, we introduce SMFNet, a generative diffusion model that predicts distributions of single-molecule chromatin accessibility profiles from DNA sequence, and PRISM, which identifies sequence features associated with individual molecular states. Together, these approaches enable us to systematically associate single molecule chromatin states at cREs with specific TF binding motifs. Experimental perturbation of these motifs redistributes the corresponding states as predicted by SMFNet, supporting their interpretation as TF-dependent configurations underlying chromatin opening. We then use controlled *in silico* motif screens to systematically dissect individual and combinatorial TF contributions to chromatin opening. Individual TFs show distinct chromatin-opening signatures, differing in accessible-region width and nucleosome-positioning precision even when associated with similar levels of chromatin accessibility. Combining motifs reveals partner- and spacing-dependent effects on chromatin accessibility, TF occupancy and nucleosome positioning. Genetic perturbation of TF binding sites using natural variation at endogenous cRE confirms that TF-specific properties shape chromatin accessibility *in vivo*. Together, our findings reveal sequence-encoded principles of combinatorial TF assembly during chromatin opening.

## Introduction

Cis-regulatory elements (cREs) control chromatin state and gene expression through the combinatorial assembly of transcription factors (TFs) on DNA. Here, TF assembly refers to their collective occupancy and activity at cREs irrespective of direct physical interactions. How DNA sequence encodes this combinatorial TF binding and activity remains poorly understood. Although large-scale *in vitro* studies have characterized the DNA-binding preferences of many mammalian TFs (1–5), motif occurrence alone does not explain occupancy *in vivo*. Most TFs occupy only a small fraction of their potential binding sites (6–8), and individual TF binding is often insufficient to establish chromatin accessibility at cREs (9). Accessible regulatory elements instead commonly contain clusters of TF binding sites (TFBSs) (10, 11), with sequence context-dependent combinatorial TF effects shaping cell-type-specific chromatin state and gene expression (10, 12–16).

TFs can influence one another through direct protein-protein interactions (17–19), DNA-mediated effects (20), or competition with nucleosomes (21, 22). These effects depend on regulatory sequence syntax, including motif identity, affinity, orientation and spacing (23–27), as well as the TF repertoire available in each cellular context (28). At endogenous elements, these features vary together, making it difficult to distinguish effects on chromatin accessibility, TF occupancy within accessible regions and nucleosome positioning.

Single-molecule genomics provides a way to resolve these joint occupancy patterns (29–36). In particular, singlemolecule footprinting (SMF) measures patterns of accessible and protected DNA along individual molecules (32, 37). In SMF, exogenous DNA methyltransferases are used to label accessible DNA, while regions protected by nucleosomes and some TFs remain unmethylated, leaving footprints of protein occupancy. Joint DNA occupancy data reveal relationships that population averages cannot uniquely determine. For example, two loci can each be individually accessible on a fraction of molecules while being accessible together on the same molecules or in mutually exclusive subsets (22, 36, 38, 39). Simultaneous measurements of chromatin accessibility on the same DNA molecule are therefore essential for distinguishing alternative configurations and quantifying their dependencies. For instance, it reveals that, at a single cRE, individual DNA molecules can engage different configurations of TF and nucleosome occupancy, each occurring at a characteristic frequency (22, 36, 38, 40). Moreover, in synthetic cREs, joint TF and nucleosome occupancy measurements have enabled thermodynamic models to link single molecule states of occupancy to transcription (41). A central challenge is therefore to understand how regulatory DNA encodes the diverse nucleosome and TF occupancy states observed across endogenous cREs.

Sequence-to-omics deep learning models offer a powerful approach to decode regulatory sequence logic from genome-wide biochemical profiling experiments (42, 43). Deep learning approaches have successfully identified sequence features that mediate regulation of TF binding (23, 44, 45), chromatin accessibility (10, 45–48), and transcription (49–56). When coupled with model interpretation methods, they have helped identify predictive motifs, their syntax and context-dependent effects (10, 23, 47, 52, 56–65). However, models trained on population-averaged measurements do not directly resolve the diversity of molecular states underlying those averages. Extending sequence modeling to single-molecule distributions would enable prediction of which sequence features support each state at the same regulatory sequence and how motif perturbations redistribute molecules across states.

To address this challenge, we introduce SMFNet, a generative diffusion model that predicts distributions of single-molecule chromatin profiles from DNA sequence, and PRISM, an interpretation method that attributes individual molecular configurations to predictive sequence features. Trained on long-read SMF data in mouse embryonic stem cells (mESCs), SMFNet captures chromatin-state frequencies and dependencies between TF footprints and nucleosome occupancy. Using controlled synthetic sequence libraries, we systematically vary TF identity, number and spacing *in silico* to determine how individual and combined TFs shape chromatin accessibility and nucleosome positioning. Individual and combinatorial motif deletions in a chromatinintegrated reporter test predicted chromatin-state redistributions, while allele-specific footprinting validates TF-specific chromatin signatures at endogenous cREs *in vivo*.

### Learning sequence-encoded chromatin-state distributions

In order to test whether DNA sequence could predict chromatin state configurations and their frequencies across single molecules, we first generated deeply sequenced genome-wide long-read SMF data in DNA methyltransferase triple-knockout (DNMT-TKO) mESCs, avoiding confounding of accessibility labeling (using exogenous CpG and GpC methyl-transferases), with endogenous CpG methylation (**Fig. 1a**). We sequenced DNA molecules continuously over 12 kilobases on average using Oxford Nanopore Technologies Sequencing (ONT) (**Extended Data Fig. E1a-b)**. These span entire cREs, simultaneously capturing multiple TF binding events and nucleosome positions. An average sequencing depth of 126x captured sufficient diversity of the chromatin states present at individual cREs to test whether their distributions can be predicted directly from DNA sequence (**Extended Data Fig. E1a**). Quantification of chromatin accessibility was highly reproducible between biological replicates and consistent with reference SMF data using bisulfite sequencing (**Extended Data Fig. E1c-d**) (22).

**Fig. 1.**
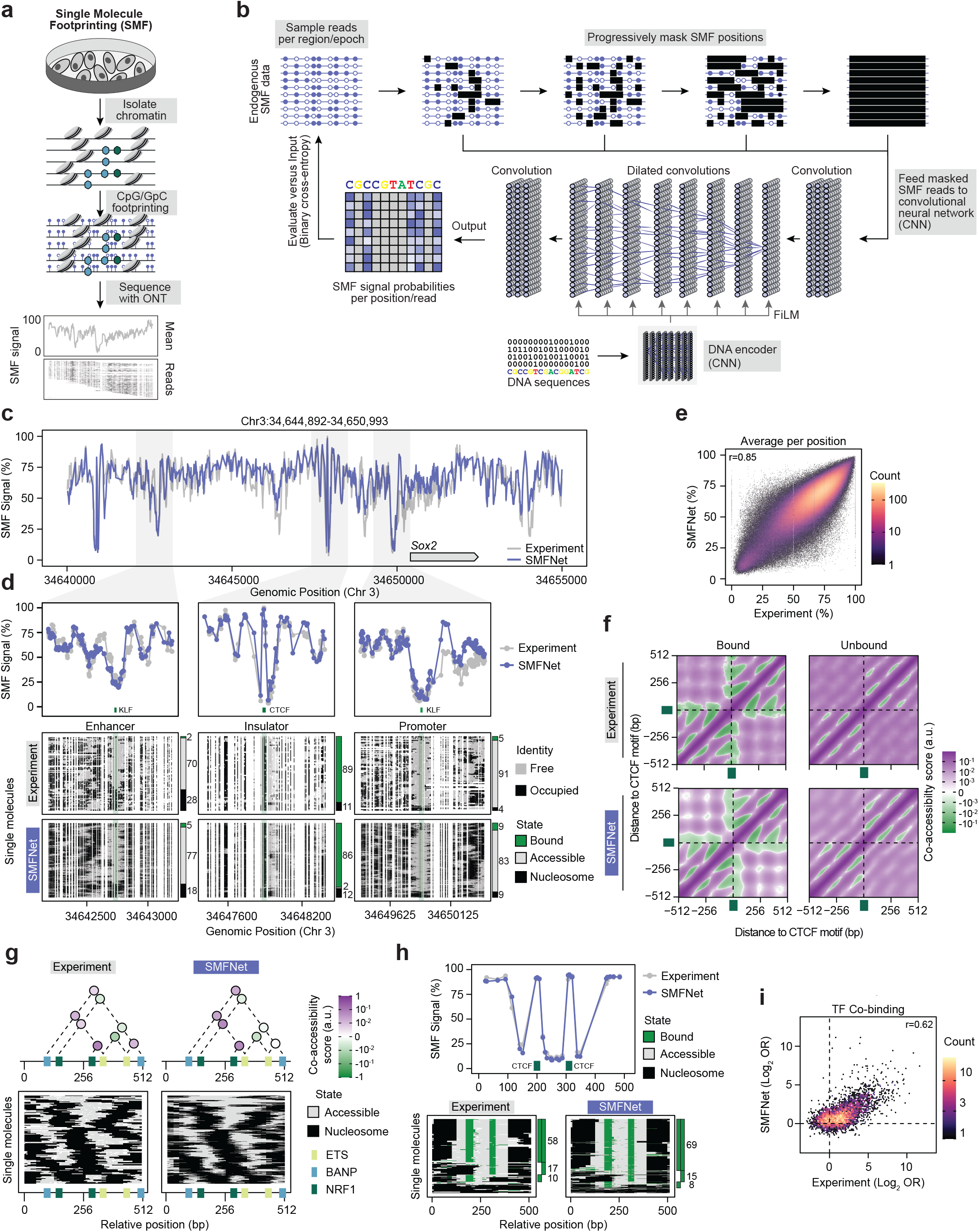
SMFNet learns single-molecule chromatin distributions and regulatory dependencies from DNA sequence. a) Schematic of the long-read SMF workflow. Oxford Nanopore SMF data were generated from DNMT triple-knockout (DNMT-TKO) mESCs in four biological replicates. **b**) Overview of the SMFNet architecture. During training, SMF reads are progressively masked by a discrete diffusion process. A convolutional neural network (CNN) is trained to reconstruct the original reads from the corrupted profiles, conditioned on the underlying DNA sequence through feature-wise linear modulation (FiLM) (71). After training, the learned denoising process samples realistic distributions of single-molecule chromatin profiles from DNA sequence alone. **c**) Average experimental (grey) and SMFNet predicted (purple) SMF profiles across the *Sox2* locus. Predictions were generated in overlapping 2048-bp windows and merged into a continuous profile. **d**) Representative enhancer (left), insulator (middle), and promoter (right) regions within the *Sox2* locus. Average SMF profiles are shown in the upper panels. Lower panels display experimental and SMFNet single-molecule chromatin profiles. Molecules are sorted according to chromatin state at a selected TF motif into bound (green), accessible (grey), and nucleosomal (black) states. The relative frequencies of each state are indicated alongside the single-molecule profiles. **e**) Correlation between experimental and SMFNet-predicted average accessibility across 2,299,914 genomic positions in held-out test regions. Only positions with >100 reads in the experimental data were included in the analysis. The Pearson’s correlation coefficient is indicated. **f)** SMFNet-generated and experimental composite co-accessibility profiles around 873 CTCF motifs from held-out test regions, stratified by reads with (Bound) or without (Unbound) a TF footprint. **g**) Co-accessibility between TF motifs at an intragenic enhancer of *Tmcc2* (Chr1:132,380,596-132,381,108). The top panels display pairwise co-accessibility scores between motif pairs, indicated above the corresponding motif locations. The bottom panels display inferred nucleosomes and accessible regions along single molecules. Only motif pairs with >100 reads overlapping both motifs were included. Annotated motifs are shown as boxes below the panels. **h**) Example CTCF-bound region (Chr1:63,180,983–63,181,495) illustrating co-binding analysis. The top panel shows average experimental (grey) and SMFNet-predicted (purple) accessibility profiles. The positions of two CTCF motifs is indicated in the plot. The lower panels display inferred state calls along single molecules including (TF-)bound (green), accessible (grey) and nucleosomal (black). The stacked bars on the right summarize the frequencies of co-binding (both motifs bound) and individual binding (only one motif bound), with the corresponding percentages indicated alongside. **i**) Scatter plot comparing co-binding in experimental and SMFNet-generated data across 4,782 TF motif pairs from held-out test regions. Only motif pairs with >100 reads overlapping both motifs and >5% bound state calls were included. The Pearson correlation coefficient is indicated in the plot.

To model these chromatin state distributions, we developed SMFNet, a generative model that uses denoising diffusion conditioned on DNA sequence to reconstruct partially hidden molecular profiles and generate new profiles through repeated reconstruction steps (66–68). During training, we mask selected methylation states (measuring chromatin accessibility) in experimental reads and train a convolutional neural network to reconstruct them using the underlying DNA sequence and the remaining visible methylation states (**Fig. 1b, Supplementary Note 1**). SMFNet comprises a DNA encoder that embeds a 2,048 bp input sequence, and a lightweight SMF predictor that uses this embedding to reconstruct the methylation state of masked sites in the central 1,024 bp of each read (**Supplementary Note 2**). To generate a read after training, SMFNet starts with all methylation states masked and fills them in over 32 steps by drawing states from the model’s predicted probabilities. Only the SMF predictor is rerun at each step, while the DNA features are computed once and reused across all steps and reads generated for a region (**Supplementary Note 3**). This allows SMFNet to learn the distribution of chromatin states encoded by each regulatory sequence rather than a single average profile. We trained SMFNet on a subset of chromosomes and evaluated its predictions on held-out test regions (**Methods**). Predicted and experimental average occupancy profiles showed strong agreement across enhancers and promoters as illustrated for the Sox2 locus (**Fig. 1c**). Across single-molecule methylation profiles, we classified protected tracts as putative TF footprints (<70 bp) or nucleosomal protection (*≥* 70 bp; **Methods**). SMFNet accurately recovered both footprint classes, including the characteristic pattern surrounding CTCF binding sites (**Fig. 1d**, middle panel). Similar accuracy was observed genome-wide when predicting average chromatin accessibility of individual cytosines (**Fig. 1e)**, or across the accessibility profiles of individual cREs **(Extended Data Fig. E1e**).

At an enhancer, an insulator and a promoter of the Sox2 locus, generated molecules reproduced the experimentally observed distributions of TF-bound, accessible and nucleosomal states (**Fig. 1d**). Consistently, SMFNet recovered the single molecule frequencies of accessible (**Extended Data Fig. E1f**), TF-bound (**Extended Data Fig. E1g**) and nucleosomal states (**Extended Data Fig. E1h**) genome-wide. Generated reads also reproduced positional correlation, capturing how chromatin accessibility states at different loci vary together across molecules, and the distribution of protected footprint lengths (**Extended Data Fig. E1i-j)**. In addition, SMFNet recapitulated aggregate SMF profiles around genomic sites containing known TF motifs (**Extended Data Fig. E1k)**. SMFNet showed comparable performance when trained on short read SMF data (**Extended Data Fig. E2a–f**). To test its performance across cell types, we trained separate models on existing single-enzyme, GpC-only SMF data from neuronal, erythroid and myoblast cell lines (69). These models predicted accessibility and dependencies between positions with accuracy comparable to that in mESCs (**Extended Data Fig. E2g-h, Supplementary Note 4**). Although SMFNet was developed for footprinting data based on cytosine methylation, it achieved comparable predictive accuracy on DNA footprinting data, which measures accessibility through adenine methylation (29, 34, 35). When trained on Fiber-seq data from the human B-lymphoblastoid GM12878 cell line (70), SMFNet captured both average accessibility profiles at individual regulatory regions and correlation between positions across single molecules (**Extended Data Fig. E3a-c, Supplementary Note 5**). Together, this demonstrates that SMFNet accurately predicts key features of single molecule chromatin state distributions at cRE.

### SMFNet predicts co-occupancy of TFs and nucleosomes on single molecules

SMF detects multiple TFs and nucleosomes on the same DNA molecule, allowing to quantify regulatory dependencies. We next asked whether SMFNet captured these dependencies by quantifying coaccessibility between genomic positions at different scales. We tiled the genome and systematically calculated how frequently two positions are simultaneously accessible or nucleosome occupied on the same molecules. At the kilobase-scale, SMFNet recapitulated the characteristic periodicity of unphased nucleosome arrays (**Extended Data Fig. E3d-e**). To assess whether SMFNet captured TF-driven changes in nucleosome organization, we stratified reads at CTCF-bound regions, where CTCF establishes strongly phased nucleosomes that locally disrupt nucleosome arrays. In both predicted and measured reads, CTCF-bound molecules showed a local disruption of nucleosome organization around its binding site (**Fig. 1f**). We next asked whether SMFNet captured co-accessibility between neighboring TF motifs within cREs. At an intragenic enhancer of *Tmcc2*, SMFNet accurately predicted both positive and negative co-accessibility associations between motifs (**Fig. 1g, Extended Data Fig. E3f**). This agreement extended genome-wide, with strong correlations between predicted and experimental co-accessibility at putative TFBSs (**Extended Data Fig. E3g**). For TFs whose binding can be resolved as short protected footprints (22), SMFNet accurately predicted TF co-occupancy profiles at individual loci (**Fig. 1h**) and their frequencies of co-occupancy genome-wide (**Fig. 1i**). Together, these results demonstrate that the dependencies between the binding of multiple TFs and nucleosome occupancy can be predicted directly from DNA sequence.

### PRISM infers TF motifs that predict single molecule chromatin states

To link predicted molecular states to the TF assemblies associated with them, we next sought to infer their sequence determinants with SMFNet. We first asked whether SMFNet recovered sequence determinants of population-averaged accessibility consistent with ChromBPNet, a convolutional neural network that predicts base-resolution bulk ATAC-seq profiles from DNA sequence (10). We trained ChromBPNet on data from the same cell type, using a peak set and chromosome splits analogous to those used for SMFNet. We quantified motif effects on accessibility by inserting 20 ChromBPNet-derived motifs into random background sequences and comparing the resulting changes in predictions between models. We then performed *in silico* mutagenesis (ISM) at 169,683 endogenous instances of these motifs, averaging mutation effects across substitutions and summing across motif positions (**Methods**). Both analyses showed concordant sequence effects between models (**Extended Data Fig. E4a–d**). Motif marginalization of cell-type-specific TF motifs revealed corresponding accessibility effects across SMFNet models trained on mESCs, neural progenitors, myoblasts and erythroleukemia cells (**Extended Data Fig. E4e**). Together, these results indicate that SMFNet recovers established motifs associated with chromatin accessibility across different cell types.

Average effects, however, do not reveal which sequence features shape individual states. We therefore developed PRISM (Per-Read In Silico Mutagenesis) to attribute individual molecular states to predictive sequence features. PRISM tests each possible single-nucleotide substitution and asks how it changes the model’s support for a chosen molecule’s methylation pattern. It samples different reconstruction orders to obtain multiple diffusion trajectories that all converge at this observed pattern. At each step, PRISM measures how the substitution changes the likelihood of the target molecule relative to the reference sequence, conditioned on the methylation states already unmasked. Averaging these changes on a logarithmic scale across reconstruction orders assigns each nucleotide a contribution score for that molecular configuration (**Extended Data Fig. E4f, Supplementary Note 6**).

At a CTCF-bound cRE, PRISM revealed markedly different attributions for different molecules (**Fig. 2a-b, Extended Data Fig. E4g**). Molecules with a single CTCF footprint were attributed specifically to the corresponding motif (**Fig. 2b, reads B and C**), molecules with two footprints to both motifs, and a nucleosome-occupied molecule showed little or no attribution to either motif (**Fig. 2b, reads A and D**). This example illustrates how PRISM can link distinct single-molecule chromatin states to specific TF-binding and nucleosome-occupancy configurations. To test whether PRISM attributions were consistent with SMFNet’s generative predictions, we introduced PRISM-selected single-base substitutions at 128 endogenous enhancers and generated profiles from the altered sequences. Higher PRISM scores corresponded to greater agreement between generated profiles and the target molecule’s methylation pattern, providing an internal check of the attribution method (**Extended Data Fig. E4h–i**). Together, these analyses support PRISM as a tool for linking individual chromatin configurations to their predictive sequence features.

**Fig. 2.**
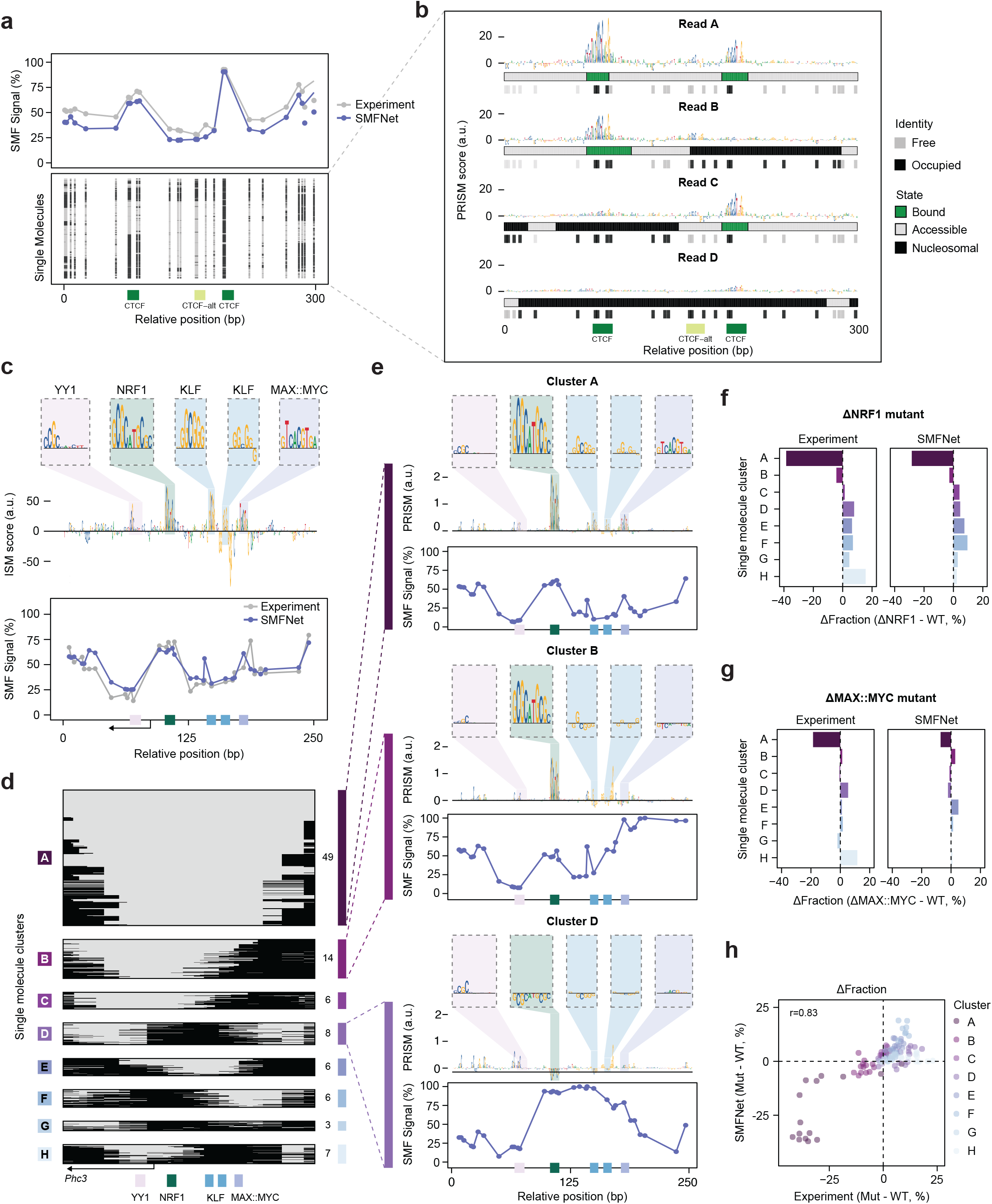
Sequence-encoded TF assemblies specify single-molecule chromatin states. **a**) SMF profiles at an example CTCF-bound CRE (Chr1:191,288,824-191,289,124). Top, average profiles from experimental SMF data and SMFNet predictions. Bottom, single-molecule chromatin profiles generated by SMFNet. **b**) PRISM (Per-Read In Silico Mutagenesis) attribution profiles for representative individual molecules. Attribution scores are shown as sequence logos, with the corresponding chromatin-state calls displayed below each read. **c**) Wildtype *Phc3* promoter analyzed in a published mCHIRA dataset (38). Top, aggregate ISM attribution scores across the sequence, with zoomed views of TF motifs. Bottom, average experimental (grey) and SMFNet generated (purple) SMF profiles. **d**) Distribution of SMFNet-generated single molecules across chromatin states at the wild-type *Phc3* promoter. Molecules were clustered according to their nucleosome-occupancy profiles using Hamming distance. Accessible regions are shown in grey and nucleosome-occupied regions in black. Values to the right indicate the relative fraction of molecules assigned to each chromatin state. **e**) Characterization of selected single molecule chromatin states at the *Phc3* promoter. Top, average PRISM attribution profiles for molecules within each cluster shown as sequence logos, with zoomed views of TF motifs. Bottom, corresponding SMFNet-generated average SMF profiles. **f-g**) Changes in single-molecule chromatin microstate frequencies following disruption of NRF1 (**f**) and MAX::MYC (**g**) motifs. Bars indicate the change in the relative abundance of each microstate between the wild-type and mutant inserts. **h**) Changes in single-molecule chromatin-state distributions across all 15 combinatorial perturbations of YY1, NRF1, MAX::MYC, and the right-hand KLF motif assayed in the mCHIRA dataset (38). The Pearson correlation coefficient is indicated.

### Identifying motif combinations associated with distinct chromatin states

We next asked whether the heterogeneous chromatin states predicted at individual cREs could be traced to distinct combinations of TF binding events. For 1,735 well-covered enhancers and promoters excluded from training, we generated 1,024 chromatin molecules and jointly clustered them with experimental reads based on their nucleosome occupancy patterns into recurrent chromatin states (**Methods**). The resulting chromatin-state frequencies agreed between generated and experimental molecules (**Extended Data Fig. E4j-k**). We then used PRISM to identify binding sites associated with specific chromatin states.

At the Phc3 promoter, ISM identified five TF motif instances associated with average predicted accessibility (**Fig. 2c**), whereas single-molecule nucleosome profiles partitioned into eight recurrent states, ranging from fully accessible (state A) to partially accessible (B–C) and largely nucleosome-occupied (D–H) configurations. These configurations closely matched those observed experimentally (**Fig. 2d, Extended Data Fig. E5a-b**). We used PRISM to assess the contribution of individual motifs to each state. The NRF1 motif contributed most strongly to states A and B, the two most frequent accessible configurations. The MAX::MYC motif contributed to state A but not state B, where it was covered by a nucleosome (**Fig. 2e, Extended Data Fig. E5c**). These state-specific sequence contributions suggest that distinct single-molecule chromatin states may be specified by different combinations of TFs binding at the same locus.

### TF assemblies specify the distribution of single– molecule chromatin states

To test these predictions, we compared SMFNet-predicted effects of individual and combinatorial motif deletions with single-molecule chromatin measurements from an integrated reporter assay. We reanalyzed our published multiplexed chromatin-integrated reporter assay (mCHIRA) dataset, which measured chromatin states on *Phc3* promoter DNA fragments integrated into the genome. TF motifs within these fragments were deleted individually or in combination, and the resulting single-molecule chromatin states were measured by targeted SMF (38). Deletion of the NRF1 motif reduced the frequency of accessible chromatin states (states A-B), and increased nucleosome-occupied states (states D-H), closely matching the SMFNet prediction (**Fig. 2f, Extended Data Fig. E5d**). Conversely, deletion of the MAX::MYC motif selectively reduced the frequency of the fully accessible state A, while modestly increasing states B and E, consistent with the different motif dependencies of states A and B and with SMFNet predictions (**Fig. 2g, Extended Data Fig. E5e**). Across both perturbations, PRISM scores at the corresponding motif in the wildtype sequence were predictive of changes in chromatin-state frequencies following motif deletion (**Extended Data Fig. E5f**). We observed similarly strong agreement across all 15 sequence variants tested (*r* = 0.83), including individual deletions of NRF1, MAX::MYC, KLF and YY1, as well as all 11 possible combinatorial motif deletions (**Fig. 2h**). Together, these results show that regulatory sequence encodes differences in chromatin-state frequencies and that motif deletions can selectively alter the states associated with distinct motif combinations.

### Individual TF motifs have distinct effects on accessibility and nucleosome positioning

At endogenous enhancers, TF identity, motif combinations and sequence context vary together, making individual motif effects difficult to isolate. We therefore used SMFNet as an *in silico* experimental system, systematically inserting and combining motifs while holding the surrounding sequence context constant. We focused on TF motifs with predicted effects on chromatin accessibility in mESCs, including pluripotency-associated (OCT4::SOX2, KLF, MAX::MYC, ESRRB, SOX2 and ZIC), and broadly expressed TFs (BANP, NFY, NRF1 and CTCF) (**Extended Data Fig. E6a, Methods**). We first assayed each motif individually to characterize its effects on chromatin accessibility. We inserted each motif into 200 randomly generated background sequences with 50% GC content, comparable to nucleotide content around endogenous mESC TFBSs (**Extended Data Fig. E6b, Methods**). We classified molecules as centrally accessible if they lacked nucleosomal protection around the sequence center, including those with short putative TF footprints. We then quantified the accessible fraction, median accessible-region width and variability in the flanking nucleosome boundaries across accessible molecules (boundary dispersion; **Fig. 3a, Methods**). Lower boundary dispersion indicates more consistently positioned nucleosomes. All measures were calculated separately for each background and then summarized as medians.

**Fig. 3.**
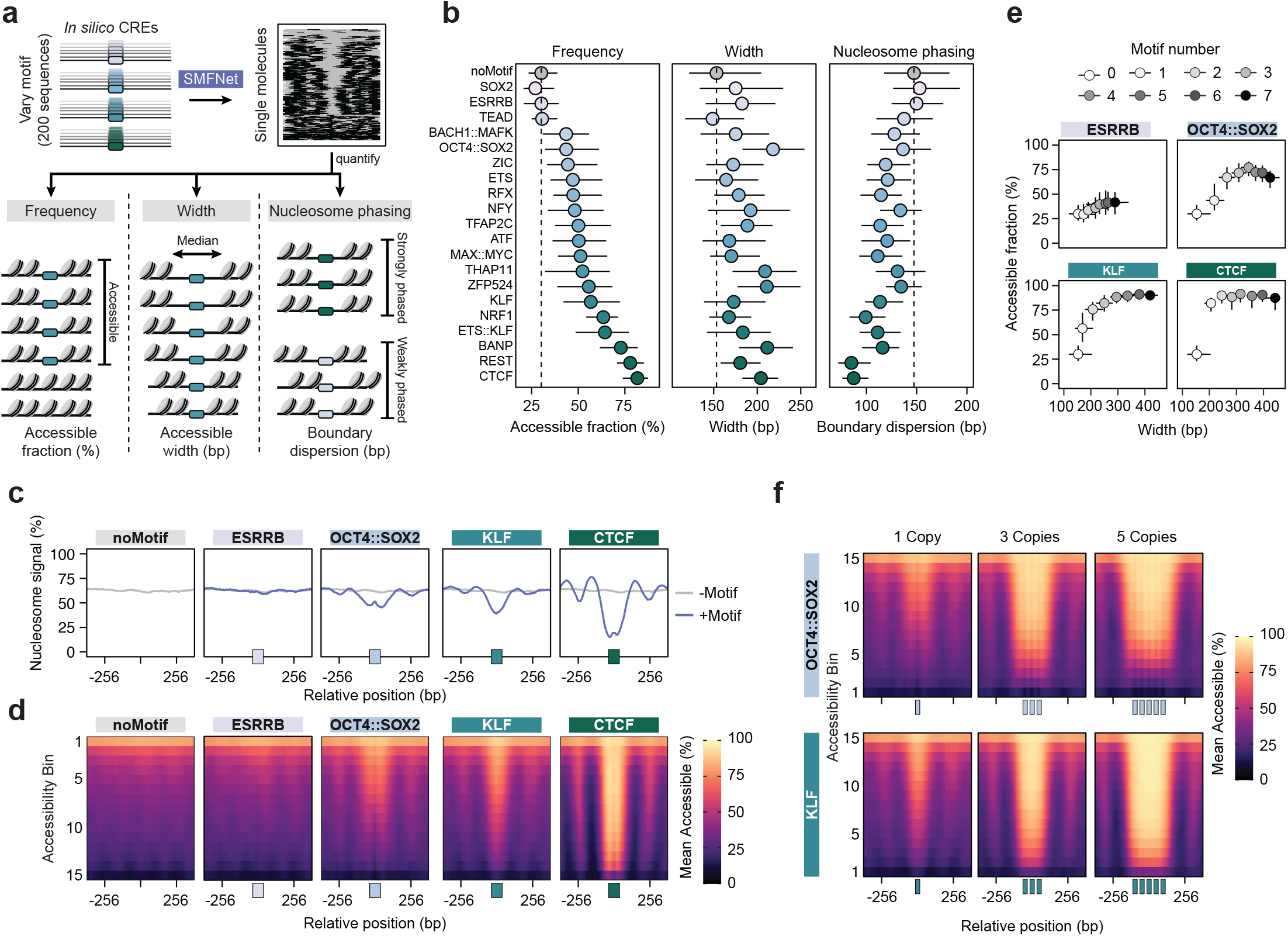
Individual TFs encode distinct chromatin-opening signatures. **a**) Schematic representation of the in silico screening approach. Single-motif insertions were sampled across 200 background sequences and quantified using three parameters: Frequency, defined as the percentage of accessible molecules. Width, defined as the median length of the accessible region. Heterogeneity, defined as the median pairwise Euclidean distance between the left and right nucleosomal boundaries of the accessible region across molecules. **b**) Frequency, width and boundary dispersion for single-motif insertions of chromatin accessibility–creating motifs. Points indicate the median across background sequences and whiskers span the interquartile range. Dashed lines indicate the corresponding median values for background sequences without motif insertion. **c**) Average nucleosome profile created following the insertion of the indicated motifs. The background regions without motif insertion are shown in grey. The position of the inserted motifs is indicated by the box below the plot. **d**) Accessibility profiles across all 200 background sequences for selected motifs. Molecules were grouped into 15 bins according to their average accessibility. Boxes below the plots indicate motif positions. **e**) Median frequency and width of chromatin opening induced by homotypic motif arrays with 50-bp centre-to-centre spacing. Whiskers denote the interquartile range. Corresponding values for sequences without motif insertion are shown in white (Motif number 0). **f**) Accessibility profiles across all 200 background sequences for one, three and five copies of OCT4::SOX2 (top) and KLF (bottom) at 50 bp spacing (center-to-center). Molecules were grouped into 15 bins according to their average accessibility. Boxes below the plots indicate motif positions.

The accessible fraction and accessible-region width varied substantially between TF motifs (**Fig. 3b**). Most motifs, including OCT4::SOX2 and KLF, increased the accessible fraction relative to controls, whereas SOX2 and ESRRB had little effect (**Fig. 3b, left panel**). For most accessibilityinducing motifs, the accessible fraction depended strongly on sequence context, including flanking GC content (**Extended Data Fig. E6c**). Motif-specific differences were also evident in average accessibility profiles and single-molecule distributions pooled across backgrounds (**Fig. 3c,d**). Motifs also differed in accessible-region width: although OCT4::SOX2 and KLF produced similar accessible fractions, OCT4::SOX2 generated substantially wider accessible regions (**Fig. 3b, center panel, Fig. 3c**,**d**). TFs also differed markedly in the precision of nucleosome positioning at the boundary of the accessible region (**Fig. 3b, right panel**). The precision of nucleosome positioning was generally correlated with the frequency of chromatin accessibility (**Extended Data Fig. E6d**). OCT4::SOX2 and NFY were notable exceptions: they positioned flanking nucleosomes less precisely than other motifs with similar frequency of chromatin accessibility (**Fig. 3b**,**d**). Together, these predictions suggest that individual TF motifs have effects on chromatin opening, shaping not only the frequency of chromatin accessibility, but also the width of accessible regions and the precision of the flanking nucleosome positioning.

We next asked how increasing the number of binding sites for the same TF alters chromatin accessibility. Adding copies of the same motif at 50-bp spacing increased both the accessible fraction and accessible-region width, but along motifspecific trajectories (**Fig. 3e, Extended Data Fig. E6e-f**). KLF arrays increased width and accessible fraction together, approaching 90% accessible molecules with five motifs. In contrast, OCT4::SOX2 increased accessible fraction and width in parallel up to three motifs, after which the accessible fraction plateaued around 75% while width continued to expand (**Fig. 3e**). These distinct trajectories were also evident in composite profiles of OCT4::SOX2 and KLF arrays (**Fig. 3f**). For most TFs, these effects were observed at spacings of 20–70 bp and weakened at larger spacings (**Extended Data Fig. E6g**). Together, these predictions suggest that each TF motif has a distinct pattern of accessibility and nucleosome positioning, and that multiple sites for the same TF act cumulatively along motif-specific trajectories.

### Motif syntax directs combinatorial chromatin opening

We next designed synthetic sequence libraries to test how motif identity and spacing shape combined effects on chromatin accessibility and nucleosome organization. Each central motif (A) was tested alone (−A−) and between two copies of a partner motif (B), across 200 matched background sequences and center-to-center spacings of 20–100 bp (BAB, Fig. 4a). We restricted the analysis to motif combinations that were frequently observed together in the training data (**Methods**). Comparing BAB with −A− measures the effect of adding the flanking motifs. To predict the accessible fraction of BAB from the component sequences, we defined a compositional accessibility baseline that combines the effects of the central motif alone (−A−) and the paired flanks (B−B), each measured relative to the empty background (−−−) (**Fig. 4a, Methods**). We estimated the combined accessibility effect of two motifs by allowing each motif to open a fraction of inaccessible molecules or close a fraction of accessible molecules. We applied these effects in both orders and averaged the results, accounting for overlapping gains and keeping expected frequencies between 0% and 100%. Width and boundary-dispersion effects were combined relative to background on a logarithmic scale (**Methods, Supplementary Note 7**). These baselines provide quantitative expectations without specifying a biological sequence of events or establishing direct TF interactions.

**Fig. 4.**
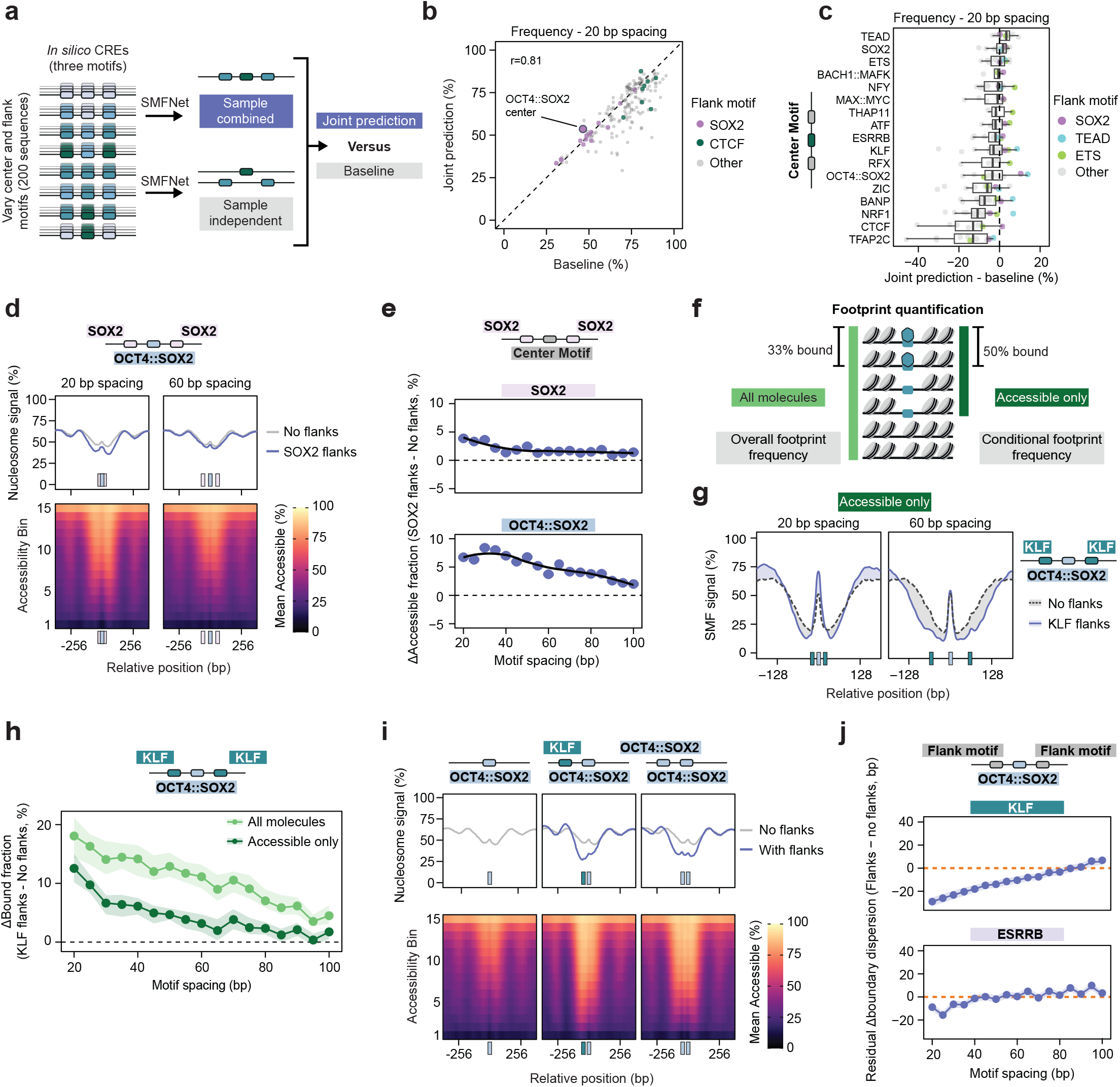
Combinatorial TF assembly shapes chromatin remodeling. **a**) Schematic of the *in silico* screening strategy. The effects of three-motif insertions on accessibility frequency, width and boundary dispersion (joint prediction) were compared with an empirical expectation derived from the corresponding individual center and flank insertions (baseline). **b**) Comparison of the frequency of accessibility predicted by the joint model and the combined-effect baseline across all motif combinations at 20 bp inter-motif spacing. Values represent medians across 200 background sequences. The Pearson correlation coefficient is indicated. **c)** Difference in frequency between the SMFNet-generated joint prediction and the corresponding baseline at 20 bp motif spacing. Each dot represents one center-flank motif combination. Thick lines indicate the median across background sequences and the boxes span the interquartile range. The whiskers denote non-outliers. **d**) Average nucleosome signal (top) and binned accessibility profiles (bottom) for OCT4::SOX2 as the center motif with SOX2 flank motifs at 20 bp and 60 bp inter-motif spacing. Molecules were grouped into 15 bins according to their average accessibility. Boxes below the plots indicate motif positions. **e**) Change in accessibility frequency following insertion of SOX2 flanks around OCT4::SOX2 and SOX2 center motifs across inter-motif spacings. The dashed line indicates the accessibility frequency of the corresponding center motif without flanks. The black lines indicate a loess smooth. **f**) Schematic of total and conditional bound fractions. Total frequency uses all analyzed molecules, while conditional frequency uses only molecules classified as accessible. The latter measures footprint prevalence within the accessible population. **g**) Smoothed average SMF signal at accessible molecules for *in silico* sequences containing a central OCT4::SOX2 motif flanked by KLF motifs at spacings of 20 bp or 60 bp. Dashed grey lines depict signal without KLF flanks. The shading highlights differences between profiles. Boxes below each panel indicate motif positions. **h**) Change in putative bound fraction following insertion of KLF flanks around a central OCT4::SOX2 motif. The data is shown for all molecules or accessible molecules only. The shaded areas indicate the 95% confidence interval. **i**) Average nucleosome signal (top) and binned accessibility profiles (bottom) for OCT4::SOX2 alone, with an additional KLF motif (middle) or with an additional OCT4::SOX2 motif (right). Flank motifs were positioned at 50 bp inter-motif spacing. Molecules were grouped into 15 bins according to their average accessibility. Boxes below the plots indicate motif positions. **j**) Frequency-adjusted change in boundary dispersion following insertion of the indicated flank motifs around OCT4::SOX2 across motif spacings. Values represent residuals from a linear model relating changes in boundary dispersion to the corresponding changes in accessibility frequency (see **Extended Data Fig. E7l**).

At 20-bp spacing, the predicted accessible fractions for combined constructs correlated with the compositional accessibility baseline (*r* = 0.81; **Fig. 4b**), but were generally lower, particularly for combinations involving motifs such as CTCF that strongly increase accessibility (**Fig. 4c**). Because the baseline accounts for overlapping accessibility gains, these deviations indicate smaller accessible fractions than expected under this combination rule. Nevertheless, flanking motifs can increase accessibility relative to the central motif alone while contributing less than the baseline predicts. Some combinations exceeded the baseline, including SOX2 motifs flanking OCT4::SOX2 (**Fig. 4b**) and specific combinations involving TEAD, SOX2 and ETS, which showed little or no chromatin-opening activity in isolation but activity in presence of a partner (**Fig. 4c**).

The accessibility gains associated with SOX2 flanks were also evident when we compared combined constructs directly with the central motif alone. SOX2 flanks increased accessibility around the central OCT4::SOX2 motif at 20-bp spacing (**Fig. 4d**). This increase declined with greater spacing (**Fig. 4e**). We observed similar effects, albeit with lower amplitude around SOX2, KLF and ESRRB centers (**Fig. 4d, Extended Data Fig. E7c**). At 60-bp spacing, the predicted accessible fractions for combined constructs correlated more strongly with the baseline (*r* = 0.97; **Extended Data Fig. E7a–b**). Similarly, the log-additive baselines likewise captured substantial variation in the width of accessibility and the precision of nucleosome positioning, with systematic differences remaining between predictions and expectations (**Extended Data Fig. E7d–g**). Thus, the effects of individual motifs broadly predict how they act in combination. However, specific motif syntax produces stronger or weaker effects on chromatin architecture than expected.

### TF partners influence binding and chromatin opening over different distances

TFs may increase the occupancy of partner TFs either indirectly through increased chromatin accessibility or directly by enhancing their binding within accessible chromatin. To disentangle these effects, we used predicted footprints as a proxy for TF binding and compared their frequency across all molecules with that among accessible molecules only (**Fig. 4f**). An increase within accessible molecules suggests that TF binding increased independently of changes in nucleosome occupancy at the cRE. We focused on synthetic sequences with a central OCT4::SOX2 motif flanked by KLF or ESRRB motifs. Neither partner produced measurable predicted footprints above background, reducing the risk of assigning flanking protection to the central motif (**Methods)**. To confirm that central footprints predicted by SMFNet reflected OCT4 binding, we trained BPNet on mESC OCT4 ChIP-seq data (23, 72) and predicted TF occupancy on the same synthetic constructs. Across spacings, changes in BPNet-predicted OCT4 counts correlated strongly with changes in total SMFNet footprint frequency for KLF flanks (*r* = 0.93), but only modestly for ESRRB flanks (*r* = 0.32; **Extended Data Fig. E7h**), supporting the ability of SMFNet to predict TF footprints. Having benchmarked these predictions, we then used SMFNet to examine TF footprints in the context of the accessibility and nucleosome occupancy states of the same molecules. At 20 bp spacing, flanking KLF motifs increased OCT4::SOX2-associated footprint frequency within accessible molecules (**Fig. 4g**). This effect was lost at 60 bp and declined sharply with increasing motif distance, approaching zero around 65 bp (**Fig. 4g–h**). In contrast, OCT4::SOX2 footprint frequency across all molecules remained elevated at larger spacings (**Fig. 4h**), suggesting that effects mediated through changes in nucleosome occupancy persist over longer distances than those acting within accessible chromatin. ESRRB produced smaller, more variable effects with a similar distance dependence (**Extended Data Fig. E7i–j**). Thus, partner effects on TF binding and chromatin accessibility operate over distinct spatial scales, consistent with a longer-range contribution of nucleosome-mediated cooperativity.

### Precision of nucleosome positioning is controlled by combinatorial TF assembly

We next asked how combinations of TFs affect the precision of nucleosome positioning. For instance, we paired OCT4::SOX2, which generates broad accessible regions with variable nucleosome boundaries, with KLF, which positions nucleosomes more consistently across molecules. Adding a KLF motif 50 bp from OCT4::SOX2 sharpened the adjacent nucleosome boundary, while the opposite boundary remained comparatively variable, producing asymmetric phasing around the cRE (**Fig. 4i**). Adding a second OCT4::SOX2 motif instead produced a weaker but more symmetric increase in phasing (**Fig. 4i**). These predictions illustrate how motif identity and placement can affect the two boundaries of an accessible region differently.

To generalize this analysis, we measured boundary dispersion, the variability of the left and right nucleosome boundaries across accessible molecules. Adding flanks generally reduced dispersion at short spacings, but this effect weakened and reversed at larger spacings (**Extended Data Fig. E7k**). Because changes in dispersion were associated with changes in accessible fraction (*r* = −0.61; **Extended Data Fig. E7l**), we fitted a linear relationship between these changes across motif combinations and spacings. We then measured each dispersion change relative to that fitted expectation. A negative residual indicates more precise positioning than expected for the accompanying change in accessibility. However, this statistical adjustment cannot determine whether the changes in nucleosome positioning and accessible fraction arise from the same molecular mechanism. For OCT4::SOX2 centers, KLF flanks produced a pronounced reduction in adjusted dispersion at short spacings that weakened progressively with distance. ESRRB and OCT4::SOX2 flanks produced smaller, oscillatory effects, including increases in adjusted dispersion at some larger spacings (**Fig. 4j, Extended Data Fig. E7m**). Oscillatory effects for ESRRB and OCT4::SOX2 were consistent with an approximately 10.5-bp helical periodicity, suggesting sensitivity to relative motif positioning along the DNA helix, reminiscent of previous observations (23). Together, these results show that motif identity and spacing jointly affect the precision of nucleosome positioning adjacent to accessible regions.

### Distinct modes of TF-driven remodeling shape chromatin at endogenous enhancers

We next asked whether the principles identified *in silico* predict TF function at endogenous enhancers. To assess motif-associated signatures at endogenous regulatory elements, we profiled mESCs derived from an F1 cross between C57BL/6J and SPRET mice, where naturally occurring single-nucleotide variants (SNVs) and small insertions or deletions (indels) alter TF motifs and are associated with allele-specific chromatin differences at thousands of loci (38, 73). In this system, the two DNA copies inherited from the parents share the same cellular environment. Sequence differences identify the parental origin of individual reads, allowing us to compare chromatin associated with the two local DNA sequences (Fig. 5a). Because each parental DNA sequence (haplotype) can carry multiple linked variants, these comparisons do not by themselves isolate the causal effect of a single motif variant. We generated deeply sequenced, long-read SMF data, achieving an average coverage of 133 kilobase-scale molecules per locus, to robustly quantify allele-specific differences in single-molecule chromatin-state frequencies (**Extended Data Fig. E8a-d**).

**Fig. 5.**
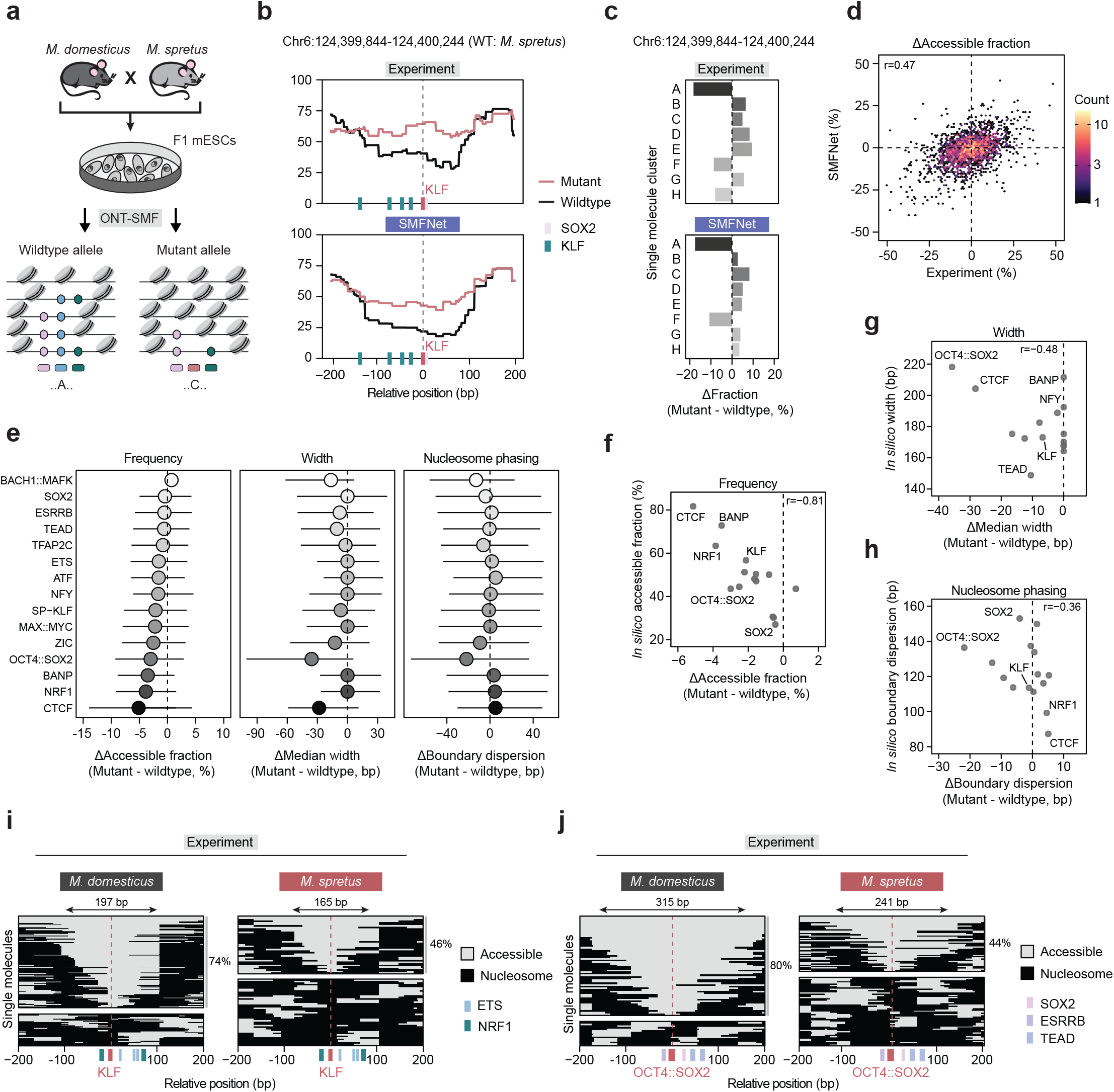
TF-specific chromatin-remodeling is recapitulated at endogenous enhancers. **a**) Schematic of the experimental workflow. F1 mESCs derived from a cross between *M. domesticus* and *M. spretus* were profiled by SMF-ONT. Changes in chromatin accessibility were quantified around naturally occurring variants disrupting annotated TF motifs. **b**) Average nucleosome profiles for wildtype and mutant alleles at a representative locus disrupting a KLF motif in experimental (top) and SMFNet-generated (bottom) data. **c**) Changes in single-molecule chromatin microstate frequencies following KLF motif disruption at the locus shown in (**b**). Bars indicate the change in the relative abundance of each microstate between the wildtype and mutant alleles. **d**) Differences in accessible fraction between *M. domesticus* and *M. spretus* alleles across 2,212 putatively deleterious TF motif variants, comparing experimental measurements with SMFNet predictions. The Pearson correlation coefficient is indicated. **e**) Changes in accessible fraction (left), median width of the accessible region (middle) and median change in boundary dispersion (right) following disruption of the indicated TF motif. The center point indicates the median and the whiskers the interquartile range. **f-h**) Concordance between median changes induced by *in silico* single motif insertion (**Fig. 3b**) and experimental motif disruption in the F1 B6xSPRET dataset for accessible fraction (**f**), median accessible width (**g**) and median boundary dispersion (**h**). Each point represents a motif tested in both settings. Pearson correlation coefficients are indicated. **i-j**) Single-molecule nucleosome profiles for *M. domesticus* and *M. spretus* alleles at representative loci disrupting a KLF (**i**, Chr1:46,905,014-46,905,414) or OCT4::SOX2 motif (**j**, Chr7:27,622,166-27,622,566). Single-molecule profiles are separated by accessibility across the variant motif. The fraction of accessible molecules is indicated adjacent to the grey bars. The median width of the accessible region is indicated above the panels. TF motifs in (**b, i-j**) are indicated by colored boxes below the plots, with the disrupted motif highlighted in red. The wildtype allele in (**b-c**) and (**e-h**) was defined as the motif variant with the higher CWM score, indicating greater similarity to the optimal TF motif.

We first tested whether SMFNet predicts chromatin differences between parental DNA copies at sites where sequence variation alters a TF motif. We matched motif instances between parental sequences by motif identity and genomic position, selected sites with the largest differences in model-derived motif scores, and compared SMFNet predictions from each parental sequence with the corresponding SMF measurements (**Methods**). At an example enhancer containing an allele-specific KLF motif disruption, SMFNet predicted differences in average accessibility (**Fig. 5b**) and in the frequencies of single-molecule chromatin states (**Fig. 5c, Extended Data Fig. E8e-f**). To compare these states, we grouped measured and model-generated molecules from both parental sequences by similarity of their nucleosome patterns, then compared the fraction of molecules in each group between alleles. This predictive agreement extended genome-wide, with SMFNet accurately capturing allele-specific changes in average accessibility (**Fig. 5d**) and, to a lesser extent, single-molecule chromatin-state frequencies (**Extended Data Fig. E8g**). Limited sampling of lowabundance states may contribute to the lower accuracy for individual state frequencies. These results show that SMFNet can predict haplotype-associated chromatin changes in complex regulatory sequences.

Stratifying allele-specific measurements by the disrupted motif revealed distinct signatures in accessible fraction, accessible-region width and boundary dispersion (**Fig. 5e**), consistent with the differences predicted from individual motif insertions (**Fig. 3b**). Disruption of CTCF, NRF1, BANP and OCT4::SOX2 produced the largest decreases in accessible fraction, whereas SOX2 and ESRRB had little effect (**Fig. 5e, left panel**). The effects on the width of accessibility also varied substantially, with OCT4::SOX2 and CTCF disruption causing the strongest reduction (**Fig. 5e, middle panel**). For most motifs associated with large increases in accessibility, nucleosome phasing was reduced on the mutated allele (**Fig. 5e, right panel**). In contrast, the loss of OCT4::SOX2 on the mutated allele increased nucleosome phasing (**Fig. 5e, right panel**), in agreement with the prediction that this TF associates with an unusual combination of high accessibility and low nucleosome phasing. These alleleassociated changes were consistent with the effects predicted *in silico* (**Fig. 3b**), but smaller at endogenous cREs (**Fig. 5e**), potentially reflecting partial motif disruption and compensatory effects of neighboring TFs on accessibility. Nevertheless, predicted TF-specific effects correlated well with the changes observed following motif disruption at endogenous enhancers (**Fig. 5f–h**). For example, OCT4::SOX2 and KLF motif variation was associated with similar changes in accessible fraction but different changes in accessibility width and nucleosome-positioning precision. These unique signatures of KLF and OCT4::SOX2 can also be observed when comparing individual enhancers where either of these motifs are disrupted. Disruption of a KLF motif reduced the accessible fraction by 28% and the width by 34 bp (**Fig. 5i, Extended Data Fig. E8h**). In contrast, disruption of an OCT4::SOX2 motif reduced the accessible fraction by a similar amount (36%), but had much higher effects on the width that changed by 74 bp (**Fig. 5j, Extended Data Fig. E8i**). Together, these comparisons support motif-associated chromatin signatures predicted *in silico* within endogenous regulatory sequences.

## Discussion

In this study, we establish a framework to learn how cRE encodes the individual and collective functions of TFs in shaping the distribution and organization of TF and nucleosome occupancy across individual DNA molecules. By coupling single-molecule footprinting with generative sequence modeling, SMFNet predicts heterogeneous chromatin states, while PRISM links these states to the TF binding sites associated with them. Genetic perturbations support these associations, showing that individual and combinatorial motif deletions selectively redistribute chromatin states. SMFNet provides an *in silico* experimental system to systematically disentangle individual and combinatorial TF functions that are otherwise obscured by the sequence complexity of endogenous cREs. Our analyses suggest that comparable levels of average chromatin accessibility can arise from markedly different underlying chromatin architectures. Individual TFs have distinct chromatin-remodeling properties that determine how frequently, how far and how precisely nucleosomes are displaced from their binding sites. These properties broadly predict their combined function, but specific TF combinations can alter their ability to open chromatin and phase nucleosomes.

Differences in the ability of TFs to engage nucleosomal DNA (5, 74–76), together with their interdependence during differentiation and reprogramming (72, 77–79), established the importance of pioneer factors in enabling other TFs to access regulatory DNA. Our predictions extend this view, suggesting that TFs influence not only how frequently chromatin is accessible but also how accessible regions are organized. For example, OCT4::SOX2 and KLF produce similar accessible fractions, but differ in width of the accessible region and the precision of nucleosome positioning. These differences may reflect TF-specific dependence on chromatin-remodeling complexes (80–82). Nucleosomebound OCT4::SOX2 can bias BAF-mediated nucleosome movement away from its binding site (83), potentially contributing to the wider accessible regions predicted for the motif. By comparison, BANP and NRF1 motifs were predicted to produce highly accessible regions with precisely positioned nucleosome boundaries. These properties may be particularly relevant at promoters (84–88), where TFdependent nucleosome positioning has been proposed to help define transcription start sites (85, 88).

The collective activity of TFs depends on how their distinct chromatin-opening properties combine. Broad agreement with our baselines suggests that combined responses are largely predictable from their individual contributions. This complements previous findings on cumulative TF contributions to transcriptional output (55) and chromatin accessibility (38). Specific combinations nevertheless exceeded these expectations, including those involving SOX2 motifs, which showed little chromatin-opening activity in isolation. Our predictions suggest that partner effects on TF binding and chromatin accessibility operate over distinct spatial scales, with binding more sensitive to motif spacing. Although the mechanism remains unclear, the flexible spacing dependence argues against a strict requirement for direct protein–protein contacts. Instead, such effects could arise through shared competition with nucleosomes (21) or TF-induced changes in DNA conformation that facilitate partner binding (89).

Sequence-based models trained on population-averaged measurements have established a powerful framework for decoding regulatory syntax, revealing motif preferences, spacing effects and context-dependent interactions governing chromatin accessibility, TF binding and gene expression (10, 23, 47–50, 52, 56, 85, 90). SMFNet builds on this foundation by predicting how regulatory sequence shapes distributions of molecular states. The same average profile can arise from different distributions of molecular configurations. and does not reveal which regulatory features co-occur on individual molecules. Predicting these joint patterns and their frequencies makes it possible to examine how sequence perturbations redistribute configurations and to assess features such as TF footprints within accessible molecules. PRISM adds sequence interpretation at the level of individual chromatin configurations by measuring how substitutions change the model’s support for each state. This extends the study of regulatory syntax from changes in aggregate signals to the molecular states and associations underlying those changes.

Our interpretation of SMFNet predictions remains constrained by the experimental assays used for training. In SMF, TF footprints can only be resolved where informative CpG or GpC sites adequately sample the protected region (37). Deaminaseand adenine methyltransferase-based approaches may offer greater positional resolution to study TF footprints in the future (29, 30, 34–36). Yet, footprint detection also depends on TF binding dynamics relative to enzyme labeling, so transient binding may leave no detectable protection. Footprint frequencies therefore report detectable protection rather than directly measuring binding affinity or residence time. Varying labeling duration can help resolve these contributions and constrain residence times with appropriate models (91). The combinatorial motif effects identified in this study are constrained by the design of the sequence libraries and the assumptions underlying the baselines. For instance, our libraries may miss some TF interactions that require precise spacings, orientations or composite sequences (20) that we did not explicitly test. In addition, exceeding our baselines does not identify the underlying mechanism of cooperativity (**Supplementary Note 7**). In the future, building explicit biophysical models based on SMFNet predictions may offer a route to connect sequence dependencies to mechanisms. SMF measurements from synthetic regulatory sequences have already been used to constrain thermodynamic and kinetic models of TF occupancy and transcription (41). SMFNet-predicted distributions could now support a similar framework across diverse sequence contexts and controlled motif perturbations to assess whether TF–nucleosome competition can account for the predicted configurations or whether additional interactions are needed.

Connecting these molecular configurations to transcription will also require resolving their dynamics. The distributions studied here do not establish whether distinct configurations represent sequential steps in chromatin opening or alternative states. Measuring changes in single molecule chromatin states upon rapid perturbations of TF function, combined with live-cell measurements of transcriptional bursting (92, 93), could help relate changes in chromatin configurations to kinetic states of transcription (94). These measurements could constrain models linking TF binding, ATP-dependent chromatin remodeling and transcription. As single-molecule assays expand to additional modalities (95– 98), their integration with generative sequence models offers a path toward understanding how regulatory sequence shapes both chromatin heterogeneity and transcriptional output (31).

## Data availability

SMF-ONT data generated from F1 B6xSpret DNMT-TKO and 159 DNMT-TKO mESCs are available in GEO under GSE350193 and GSE350195.

Trained SMFNet models for DNMT-TKO mESC SMFONT and GM12878 Fiber-seq data are available on Hugging Face (https://huggingface.co/collections/kundajelab/smfnet). Additional models, processed training inputs, sampled reads and DNA sequences used for sampling are available on Zenodo (DOI: 10.5281/zenodo.23407172).

## Code availability

The SMFNet package and code to reproduce the manuscript analyses are available at the following repositories.

SMFNet package https://github.com/kundajelab/SMFNet/

Manuscript analysis code https://github.com/kundajelab/SMFNet_paper_analysis_code

## Supplementary data

*Supplementary Table 1*: Generated sequencing data and published datasets used in this study

*Supplementary Table 2*: SMFNet model training and benchmarking (Related to Fig. 1 and Extended Data Fig. E1-4)

*Supplementary Table 3*: mCHIRA cluster and PRISM analyses (Related to Fig. 2 and Extended Data Fig. E4-5)

*Supplementary Table 4*: *In silico* syntax analyses (Related to Fig. 3-4 and Extended Data Fig. E6-7)

*Supplementary Table 5*: F1 B6xSpret SMF-ONT analysis (Related to Fig. 5 and Extended Data Fig. E8)

## Acknowledgments

We thank Michael Dorrity, Brian Trippe, Andreas Bally, Michela Palamin and Bram Verhagen for feedback on the manuscript. We thank Tino Hochepied and Claude Libert for sharing the B6xSpret mESC line.

We acknowledge MODIS and GeneCore EMBL for support in sequencing data generation and management respectively. We thank PepCore for the production of recombinant methyl-transferases. We thank EMBL IT Services and Stanford Research Computing for support with high-performance computing infrastructure and cluster management.

Research in the laboratory of A.R.K. is supported by EMBL core funding, the Deutsche Forschungsgemeinschaft (KR 5247/1-3), and the ERC (TFCoop-101125530). T.S. is supported by the EMBL-Stanford Life Science Alliance. N.D. is supported by the Stanford Graduate Fellowship in Science & Engineering.

## Contributions

T.S, A.K. & A.R.K conceptualized the project, T.S. & N.D. developed SMFNet and performed benchmarking, N.D. developed PRISM, implemented Fiberseq and GpC-only SMFNet, and established the SMFNet Github repository, T.S. performed co-dependency analysis and model interpretation with help of N.D., T.S. performed *in silico* screens and F1 SMF data analysis, Y.B. generated long-range SMF data, R.K. generated F1 SMF data with help of Y.B. and T.S., A.R.K & T.S. wrote the manuscript with help of A.K. & N.D. A.R.K. and A.K. acquired the funding. A.R.K and A.K. supervised the work. All authors discussed the results and approved the manuscript.

---

**Extended Data Fig E1.**
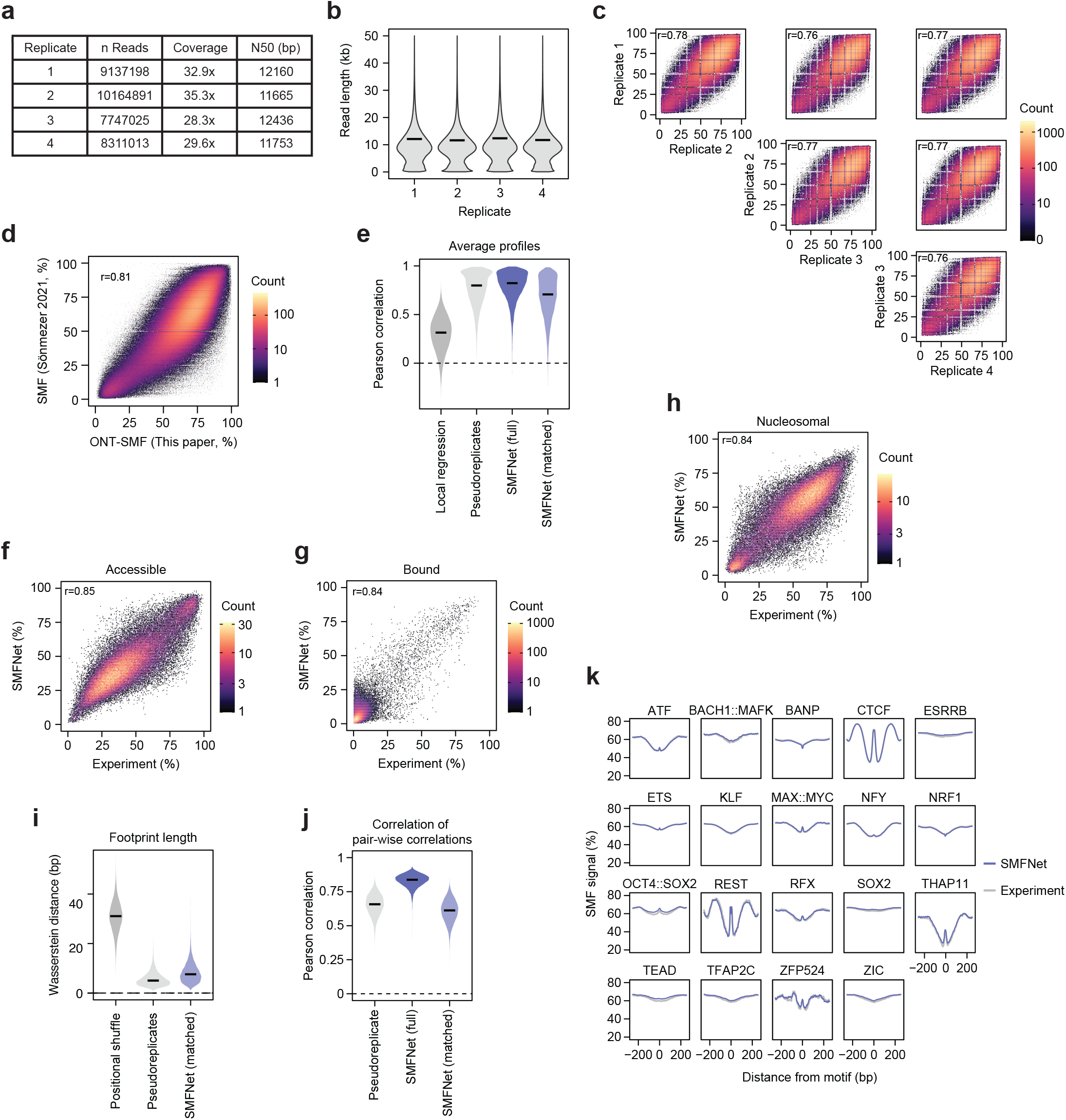
Quality control of SMF-ONT data and benchmarking of SMFNet predictions. **a**) Summary statistics of long-read SMF sequencing data from DNMT-TKO mESCs, shown separately for each biological replicate. **b**) Read-length distributions of long-read SMF data from TKO mESCs, shown by replicate. Black bars indicate the N50 read length. **c**) Average SMF signal at individual SMF positions across biological replicates. Only positions covered by at least 20 reads in all replicates were included. Pearson correlation coefficients are indicated in the plots. **d**) Average SMF signal per position in the newly generated long-read SMF dataset compared with short-read Illumina SMF data generated in the same cell type (22). **e**) Pearson correlation of average SMF profiles across 13,716 held-out test regions for a local regression baseline, experimental pseudoreplicates, and SMFNet predictions using either all 1,024 generated reads (full) or a subset matched in size to one half of the experimental reads (matched; see Methods). The baseline uses logistic regression trained to predict each site’s mean SMF signal from its ±5 bp sequence context, with separate models for CpG, GpC, and GCG sites. Black bars indicate the median. **f-h**) Fraction of accessible (**f**), bound (**g**), and nucleosomal (**h**) state calls at 74,086 candidate TF motifs in held-out test regions. Pearson correlation coefficients are indicated in the plots. **i-j**) Wasserstein distance between protected-region length distributions (**i**) and Pearson correlation between positional correlation matrices (**j**) across 13,716 held-out test regions. Both panels show experimental pseudoreplicates and size-matched SMFNet predictions (matched; see Methods). (**i**) additionally includes position-shuffled controls, and (**j**) includes predictions using all 1,024 generated reads (full). Black bars indicate the median. **k**) Composite SMF profiles centered on candidate TF motifs in SMFNet-generated (purple) and experimental (grey) data. Only motifs from held-out test regions were included

**Extended Data Fig E2.**
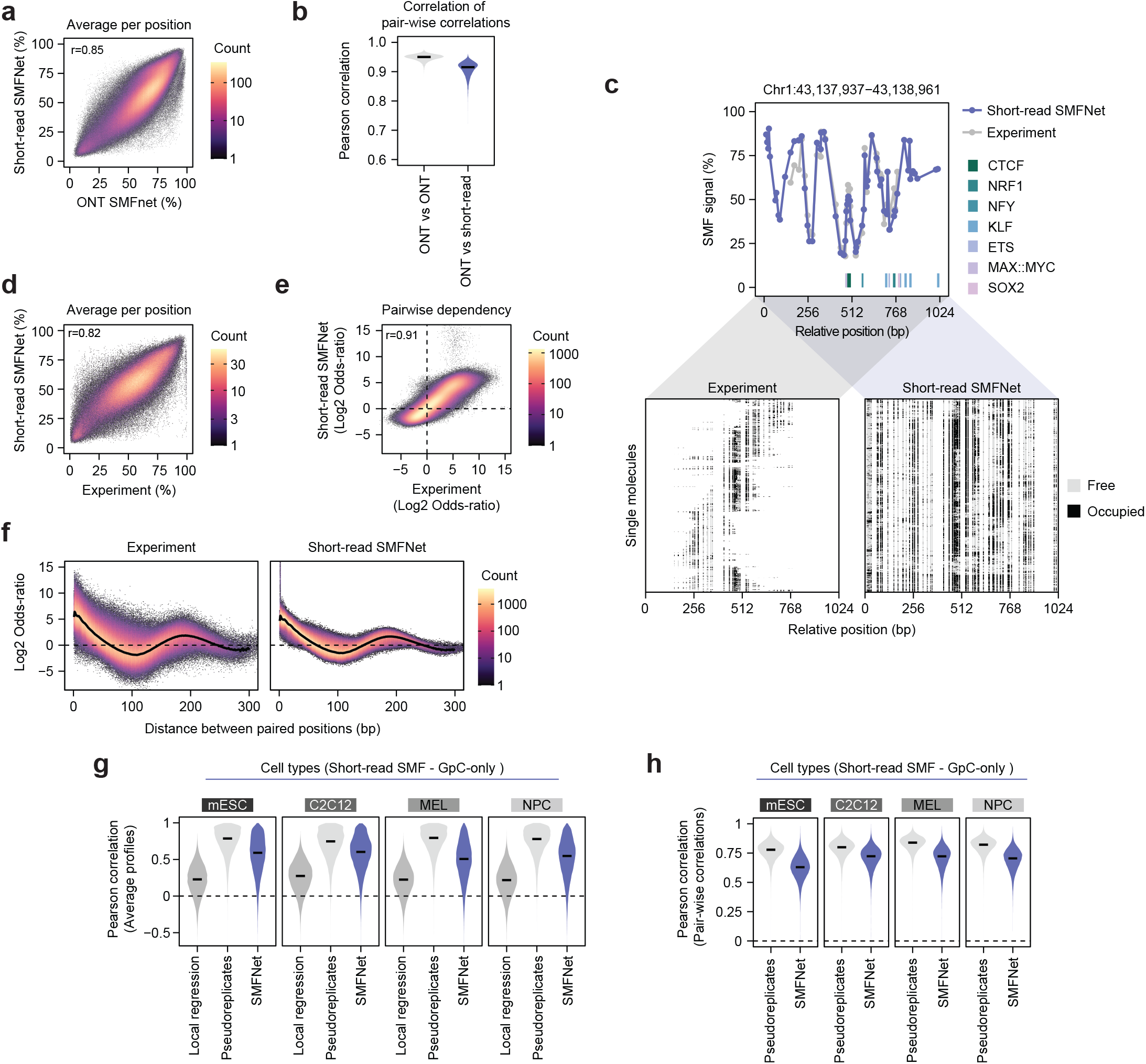
SMFNet performance across sequencing platforms and cell types. **a**) Average SMF signal at 2,307,729 individual SMF positions across held-out test regions predicted by the long-read ONT and short-read Illumina SMFNet models. The Pearson correlation coefficient is indicated. **b**) Pearson correlation between pair-wise correlation matrices across 15,505 held-out test regions. Results are shown for two non-overlapping subsets of ONT SMFNet predictions (ONT vs ONT) and short-read SMFNet versus ONT SMFNet predictions (ONT vs short-read). The black bars denote the median. **c**) Example region displaying average (top) and single molecule profiles (bottom) of experimental and short-read SMFNet-generated data. Methylated (accessible) positions are depicted in grey, while unmethylated (occupied) positions are colored in black. Positions of putative TFBSs are annotated below the plot. **d**) Average SMF signal at 565,726 individual SMF positions across held-out test regions in experimental and short-read SMFNet-generated data. The Pearson correlation coefficient is indicated. **e**) Comparison of pairwise dependencies across 5,945,552 SMF position pairs in experimental and short-read SMFNet-generated data, quantified as log2 odds ratio. The Pearson correlation coefficient is indicated. **f**) As in **(e)**, stratified by the distance between paired positions. The thick black line indicates the generalized additive model (GAM) fit. **g–h**) Pearson correlation of average SMF profiles (**g**) and positional correlation matrices (**h**) across 28,360–30,693 held-out test regions using short-read, GpC-only SMF data (69) from embryonic stem cells (mESC), myoblasts (C2C12), erythroleukemia cells (MEL), and neural progenitor cells (NPC). Experimental pseudoreplicates and SMFNet predictions based on 1,024 generated reads are shown in both panels. (**g**) additionally includes a local logistic regression baseline trained to predict each GpC site’s mean SMF signal from its ±5 bp sequence context. Black bars indicate the median.

**Extended Data Fig E3.**
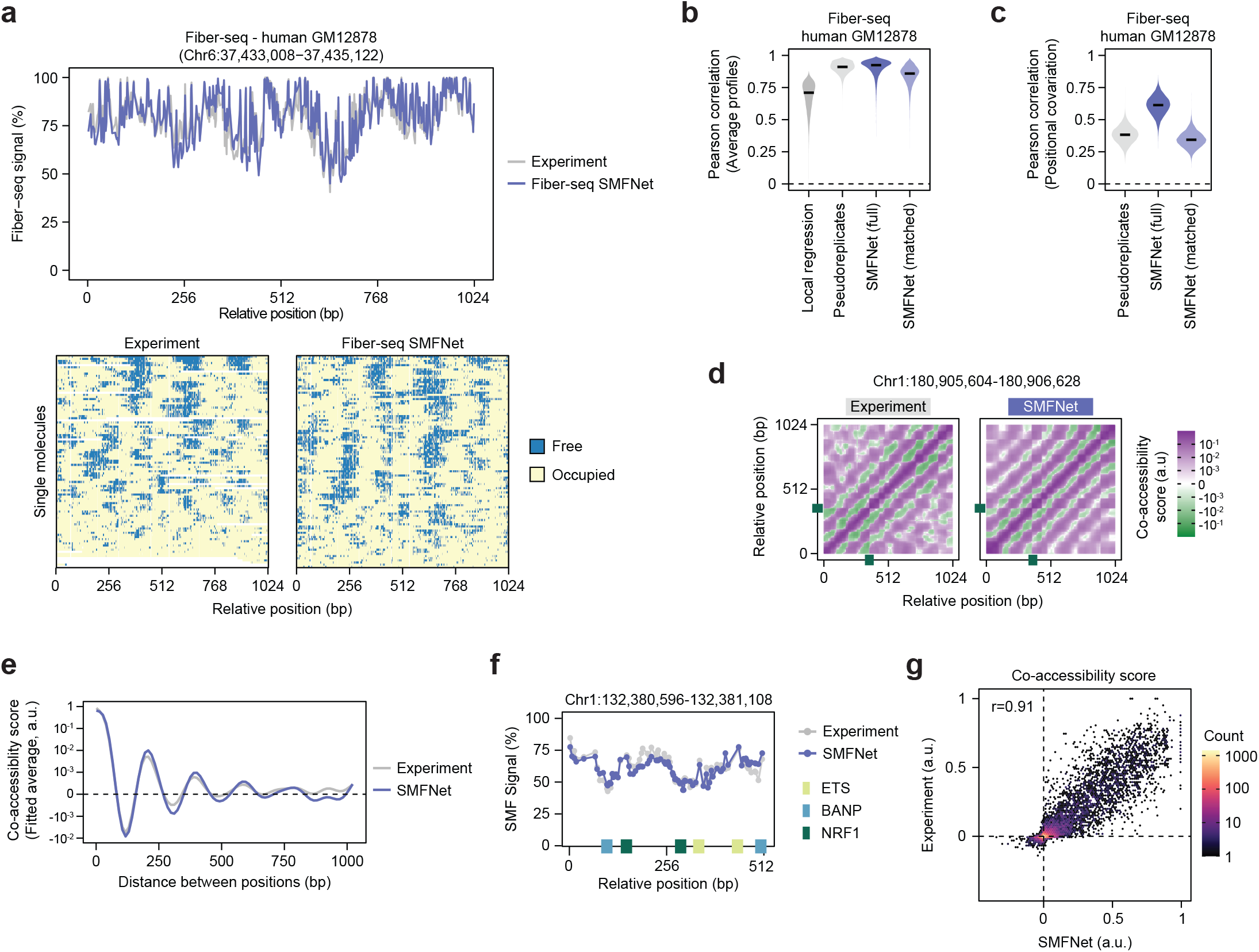
Fiber-seq benchmarking and prediction of chromatin co-accessibility. **a)** Example region displaying average (top) and single molecule profiles (bottom) of experimental and SMFNet-generated Fiber-seq data in human GM12878 cells (70). Methylated (accessible) positions are depicted in blue, while unmethylated (occupied) positions are colored in pale yellow. The single molecule panels show 100 randomly sampled reads. **b-c)** Pearson correlation of average Fiber-seq profiles (**b**) and positional correlation matrices (**c**) across 17,384 held-out test regions in human GM12878 cells. Results are shown for experimental pseudoreplicates and SMFNet predictions using either all 1,024 generated reads (full) or a subset downsampled to match the experimental read depth (matched). Black bars indicate the median. **d**) Unseen example region showing position-by-position co-accessibility matrices for experimental and SMFNet-generated reads. The green box indicates a CTCF motif. **e**) GAM fit of co-accessibility as a function of genomic distance for 23,572,969 pairs of SMF positions from unseen test regions. **f)** Average SMF profiles from experimental and SMFNet-generated reads for an example region used to display co-accessibility analysis. Annotated TF motifs are shown as boxes. **g**) Comparison of co-accessibility scores between experimental and SMFNet-generated reads for 10,297 TFBS pairs from unseen test regions. Pearson correlation is indicated.

**Extended Data Fig E4.**
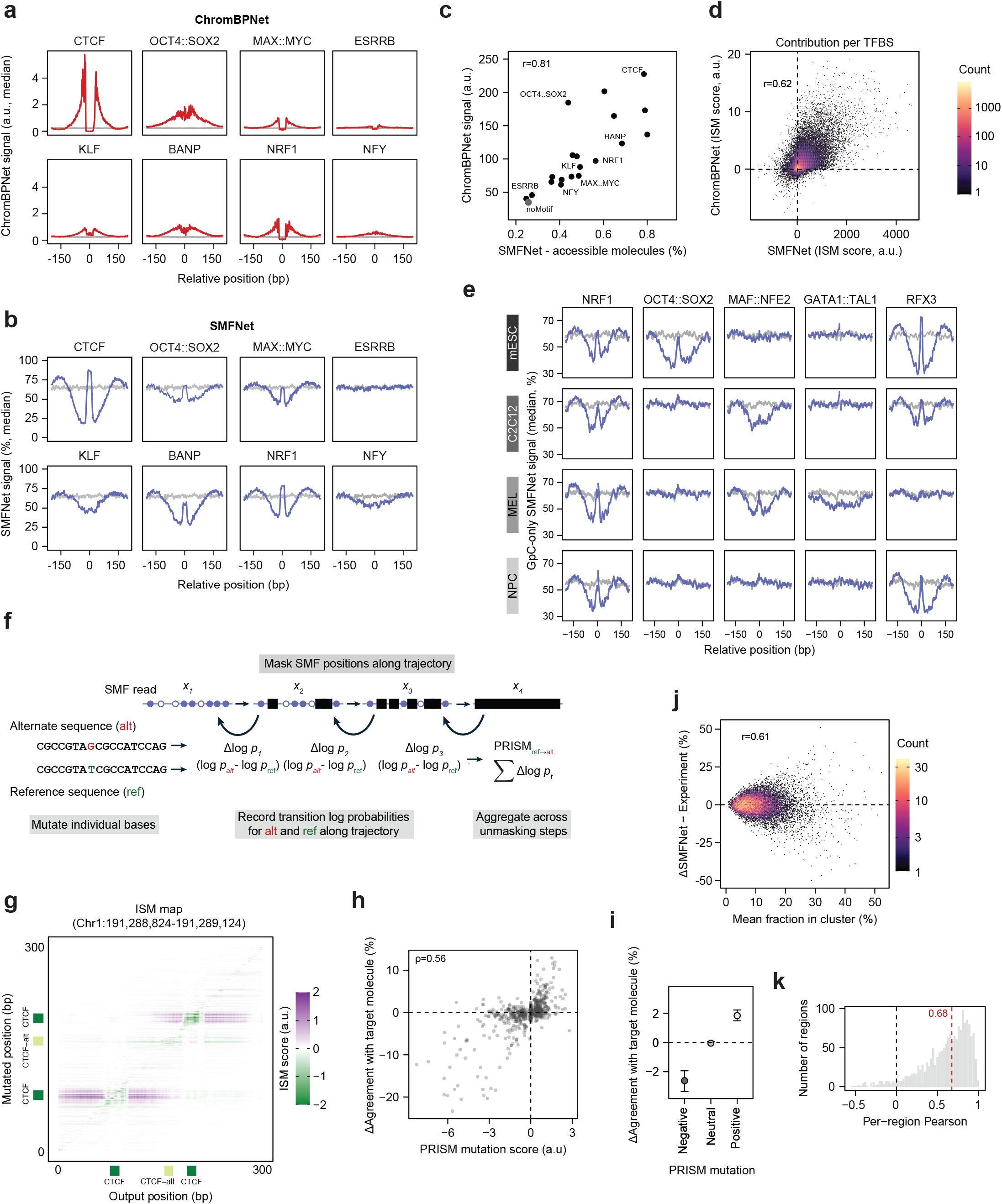
Benchmarking sequence effects and chromatin-state interpretation with PRISM. **a-b**) Composite plots showing median ChromBPNet (**a**) and SMFNet (**b**) signal following insertion of selected TF motifs into 200 randomly generated background sequences. The median signal without motif insertion is shown in grey. **c**) Comparison of motif-induced accessibility predicted by ChromBPNet and SMFNet, calculated as the median across 200 random background sequences. Background sequences without motif insertion are shown in grey. The Pearson correlation and selected TF motifs are indicated. **d**) Comparison of ISM scores from SMFNet and ChromBPNet across 169,683 putative TFBSs in unseen test regions. Per-TFBS scores were calculated by summing base-pair-level ISM scores across each motif. Pearson correlation is indicated. **e)** Composite plots showing median short-read, GpC-only SMFNet signal following insertion of selected TF motifs into 200 randomly generated background sequences for models trained in mouse embryonic stem cells (mESC), neural progenitor cells (NPC), myoblasts (C2C12), and erythroleukemia cells (MEL). The median signal without motif insertion is shown in grey. **f)** Schematic of the PRISM algorithm for a single-base mutation. At each unmasking step, transition log probabilities are calculated for the reference and alternative sequences, and their differences are summed to obtain the PRISM score. Repeating this across all positions generates a sequence contribution profile for an individual SMF molecule. **g)** ISM map showing position-by-position sequence contributions across an example region. Annotated TF motifs are indicated by boxes. Scores represent the mean change in predicted signal in logit space across all alternative single-base substitutions at each position. **h**) Correlation between PRISM-predicted mutation effects and changes in methylation agreement with the target molecule, calculated SMFNet-generated reads across 128 endogenous enhancers. Two experimental molecules per enhancer were randomly selected as target profiles. Mutations with the most positive, most negative, and closest-to-zero PRISM scores were evaluated for each target. The Spearman correlation coefficient is indicated. **i**) Mean change in methylation agreement with target molecules for the mutations in (**h**), grouped by predicted mutation effect. Error bars indicate 95% confidence intervals estimated from 2,000 bootstrap replicates. **j**) Difference in cluster frequencies between SMFNet-generated and experimental reads across 1,735 candidate CREs from unseen test regions, plotted against the mean frequency across both datasets. Pearson correlation is indicated. **k**) Region-wise Pearson correlation comparing cluster-frequency distributions between experimental and SMFNet-generated reads across 1,735 candidate CREs from unseen test regions.

**Extended Data Fig E5.**
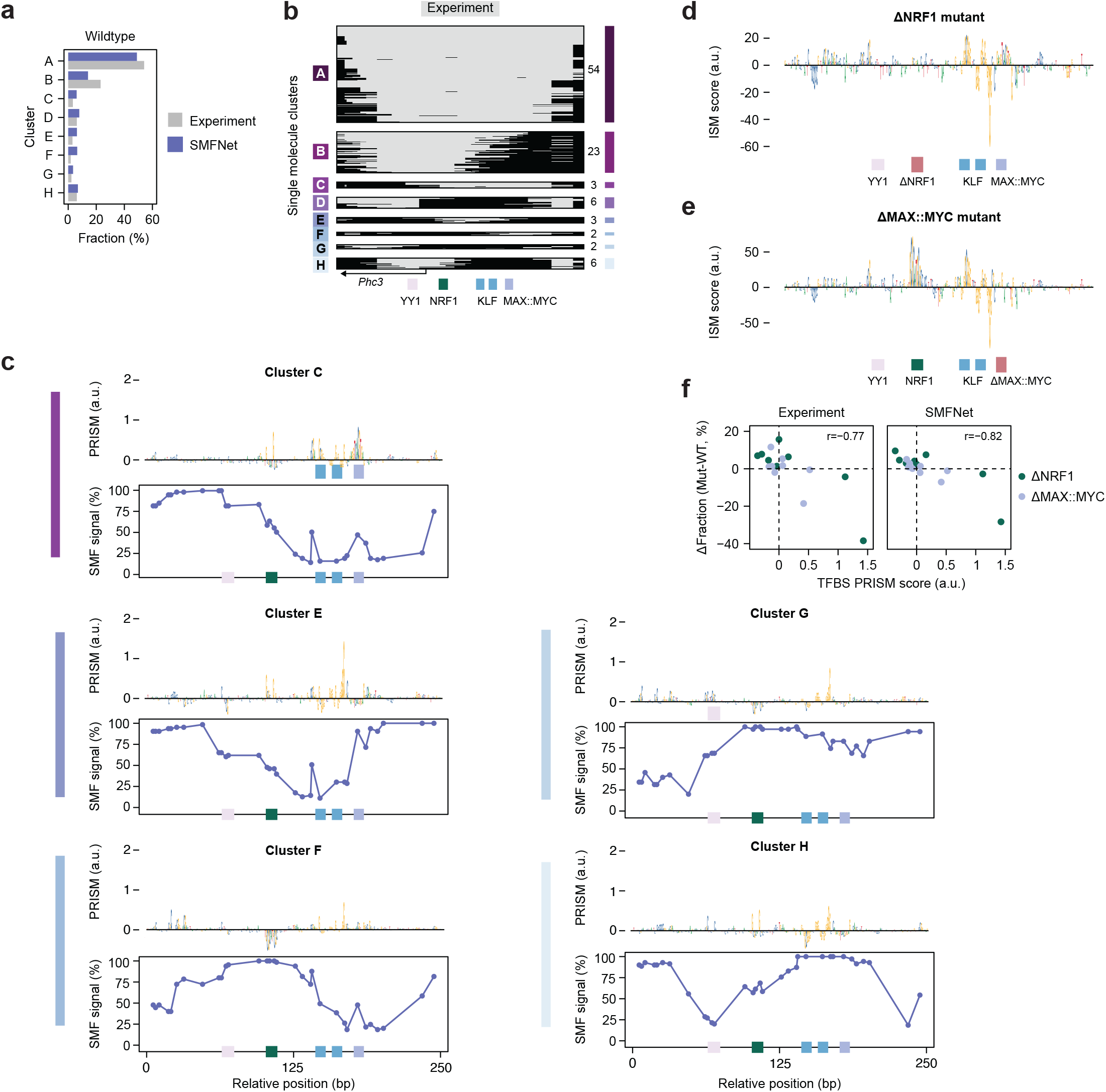
Sequence determinants of chromatin states at the *Phc3* promoter. **a**) Relative single-molecule chromatin state frequencies of experimental and SMFNet-generated molecules in the wild-type *Phc3* promoter. **b**) Distribution of experimental single molecules across chromatin states at the wild-type *Phc3* promoter. Molecules were clustered according to their nucleosome-occupancy profiles using Hamming distance. Accessible regions are shown in grey and nucleosome-occupied regions in black. Values to the right indicate the relative fraction of molecules assigned to each chromatin state. **c**) Characterization of selected single molecule chromatin states at the *Phc3* promoter. Top, average PRISM attribution profiles for molecules within each cluster shown as sequence logos. Bottom, corresponding SMFNet-generated average SMF profiles. **d-e**) Aggregate ISM attribution scores for the *Phc3* promoter following disruption of the NRF1 (**d**) and MAX::MYC (**e**). **f**) Correlation between wildtype PRISM scores for the NRF1 and MAX::MYC motifs across chromatin-state clusters and the corresponding changes in cluster frequency following motif disruption, as predicted by SMFNet and observed experimentally. TFBS PRISM scores were quantified as the mean score across the core motif. Pearson correlation coefficients are indicated.

**Extended Data Fig E6.**
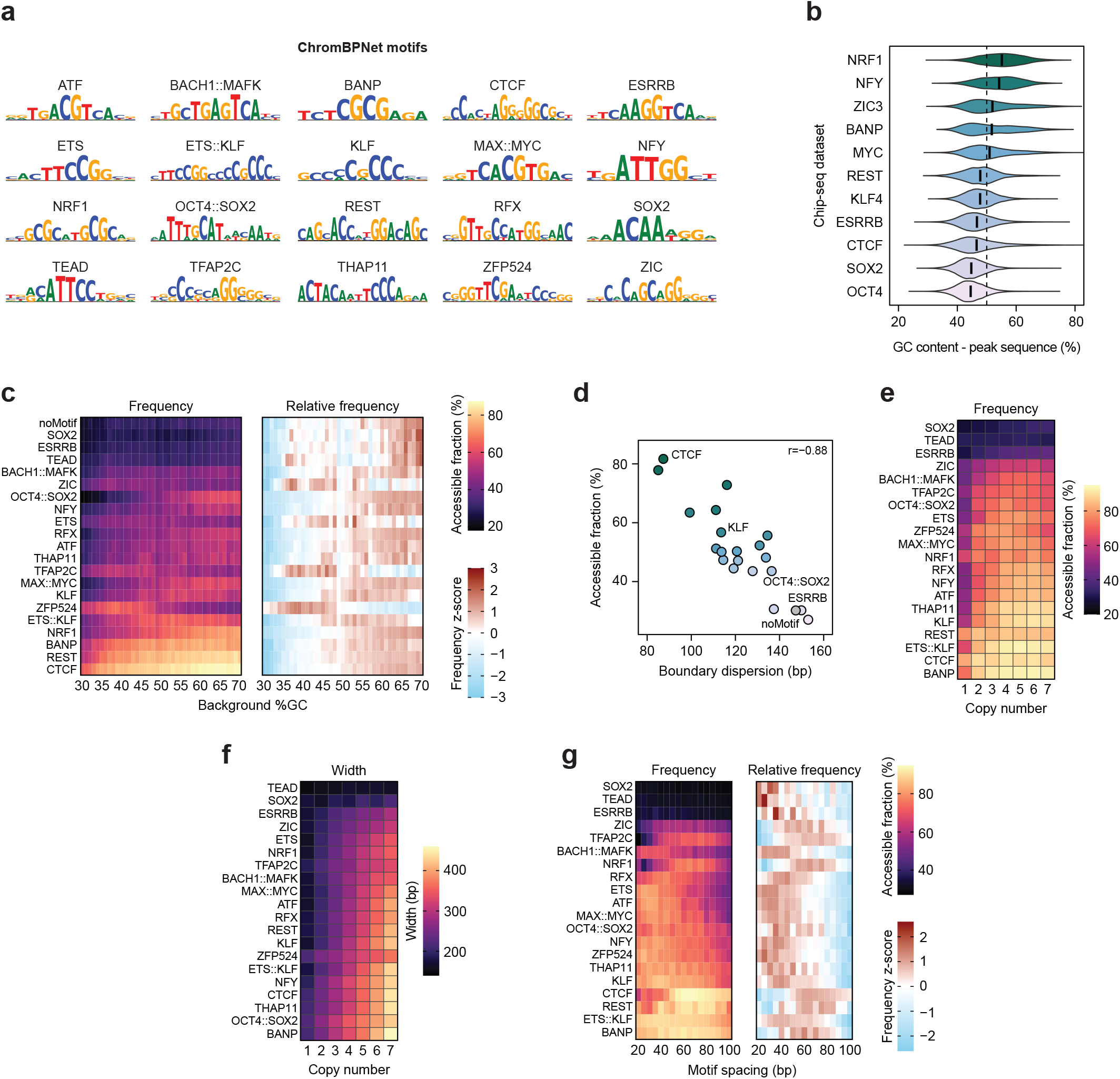
Influence of sequence context, motif number and spacing on predicted chromatin opening. **a**) Contribution weight matrices identified by TF-MoDISco from a ChromBPNet model trained on DNMT triple-knockout mESCs (10, 65, 99) and used for the *in silico* motif insertions. **b**) GC content of genomic sequences surrounding mESC ChIP-seq peak summits (±500 bp) for the indicated TFs. **c**) Absolute (left) and z-score normalized (right) median accessible fraction following single motif insertion across background sequences of varying GC content. 200 background sequences were assayed per GC percentage bin. **d**) Relationship between accessible fraction and boundary dispersion. Values represent the median across 200 background sequences. Pearson correlation coefficients and selected motifs are indicated. **e-f)** Median accessible fraction (**e**) and width (**f**) induced by homotypic motif insertions with a motif spacing of 50 bp (center-to-center). **g**) Absolute (left) and z-score normalized (right) median accessible fraction for homotypic triple motif insertions using the indicated motif spacings (center-to-center).

**Extended Data Fig E7.**
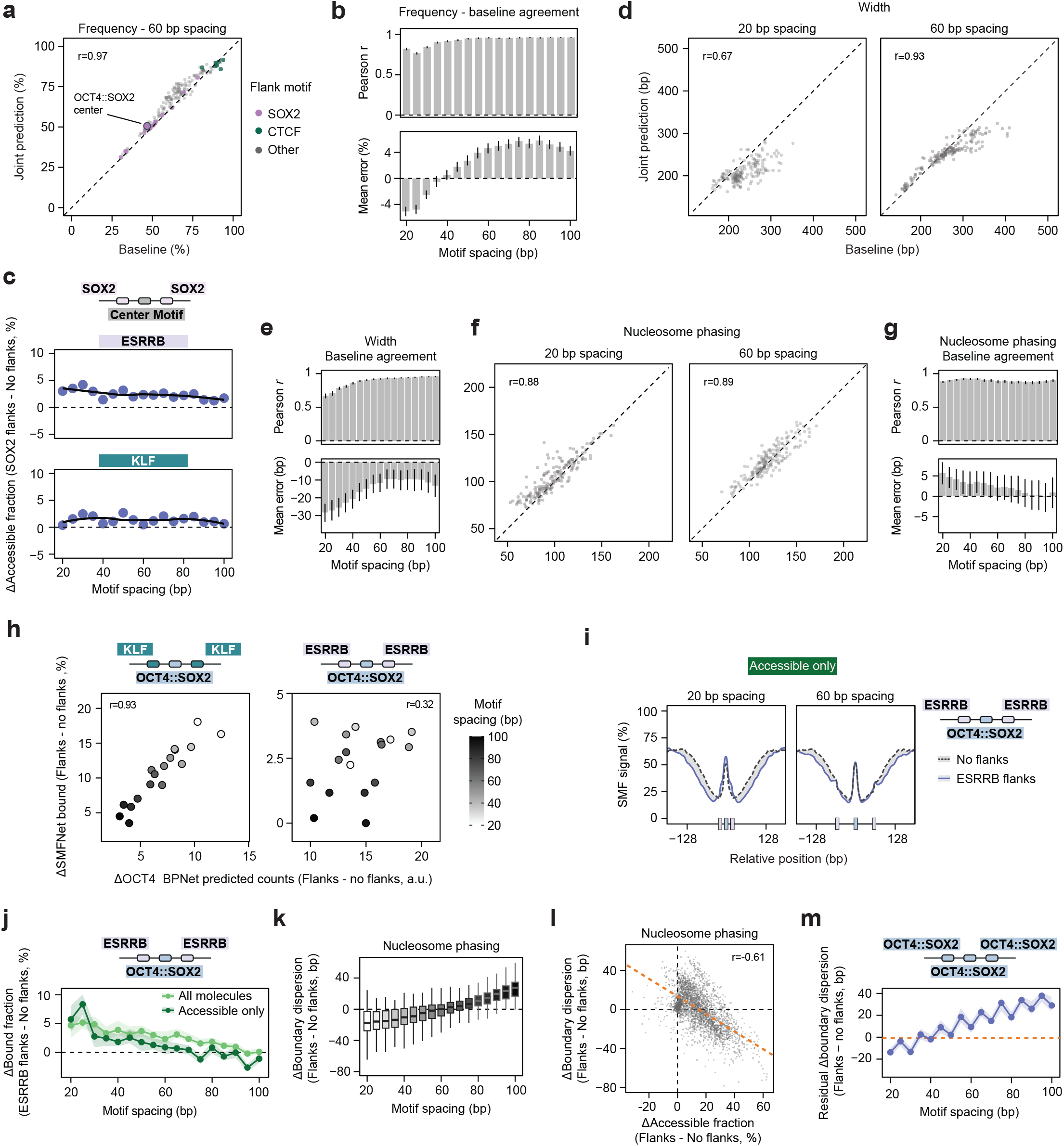
Combinatorial motif effects on accessibility, TF occupancy and nucleosome positioning. **a**) Predicted accessible fractions for combined constructs and expectations from the compositional accessibility baseline at 60-bp spacing. Values are medians across 200 backgrounds. The dashed line indicates equality and Pearson r is shown. **b**) Pearson correlation (top) and mean signed difference (bottom) between joint predictions and baseline expectations for accessible fraction across spacings. Black intervals indicate 95% confidence intervals from background bootstrap resampling. **c**) Change in accessible fraction upon adding SOX2 flanks around ESRRB or KLF centers. The dashed line denotes no change relative to center-only constructs. The black curves show LOESS fits. **d–g**) Comparison of joint predictions and log-additive baseline expectations for accessible-region width **(d–e)** and boundary dispersion (**f–g**) across motif combinations at the indicated inter-motif spacings. Values are medians across 200 background sequences. Scatter plots (**d, f**) show Pearson correlation coefficients. Summary plots (**e, g)** show Pearson correlation (top) and mean error (bottom), with black lines indicating 95% confidence intervals. **h**) Correlation between changes in SMFNet-predicted bound frequency at a central OCT4::SOX2 motif and BPNet-predicted OCT4 ChIP counts upon adding flanking KLF (left) or ESRRB (right) motifs at the indicated inter-motif spacings. Changes were calculated relative to the same sequences without flanking motifs, then summarized as medians across 200 *in silico* sequences. Pearson correlation coefficients are indicated. **i**) Smoothed average SMF signal at accessible molecules for *in silico* sequences containing a central OCT4::SOX2 motif flanked by ESRRB motifs at spacings of 20 bp (top) or 60 bp (bottom). Dashed grey lines depict signal without ESRRB flanks. The shading highlights differences between profiles. Boxes below each panel indicate motif positions. **j**) Change in putative bound fraction following insertion of ESRRB flanks around a central OCT4::SOX2 motif. The data is shown for all molecules or accessible molecules only. The shaded areas indicate the 95% confidence interval. **k**) Change in boundary dispersion following insertion of flank motifs across motif combinations at the indicated inter-motif spacings. Thick lines indicate the median across background sequences and the boxes span the interquartile range. The whiskers denote non-outliers. **l**) Relationship between changes in boundary dispersion and accessible fraction upon adding flanks across retained motif combinations and spacings of 20–100 bp. The orange line shows the fitted linear relationship. The Pearson correlation coefficient is indicated. **m**) Frequency-adjusted change in boundary dispersion following insertion of OCT4::SOX2 flanks around OCT4::SOX2 across inter-motif spacings. Values represent residuals from a linear model relating changes in boundary dispersion to the corresponding changes in accessibility frequency (see **l**).

**Extended Data Fig E8.**
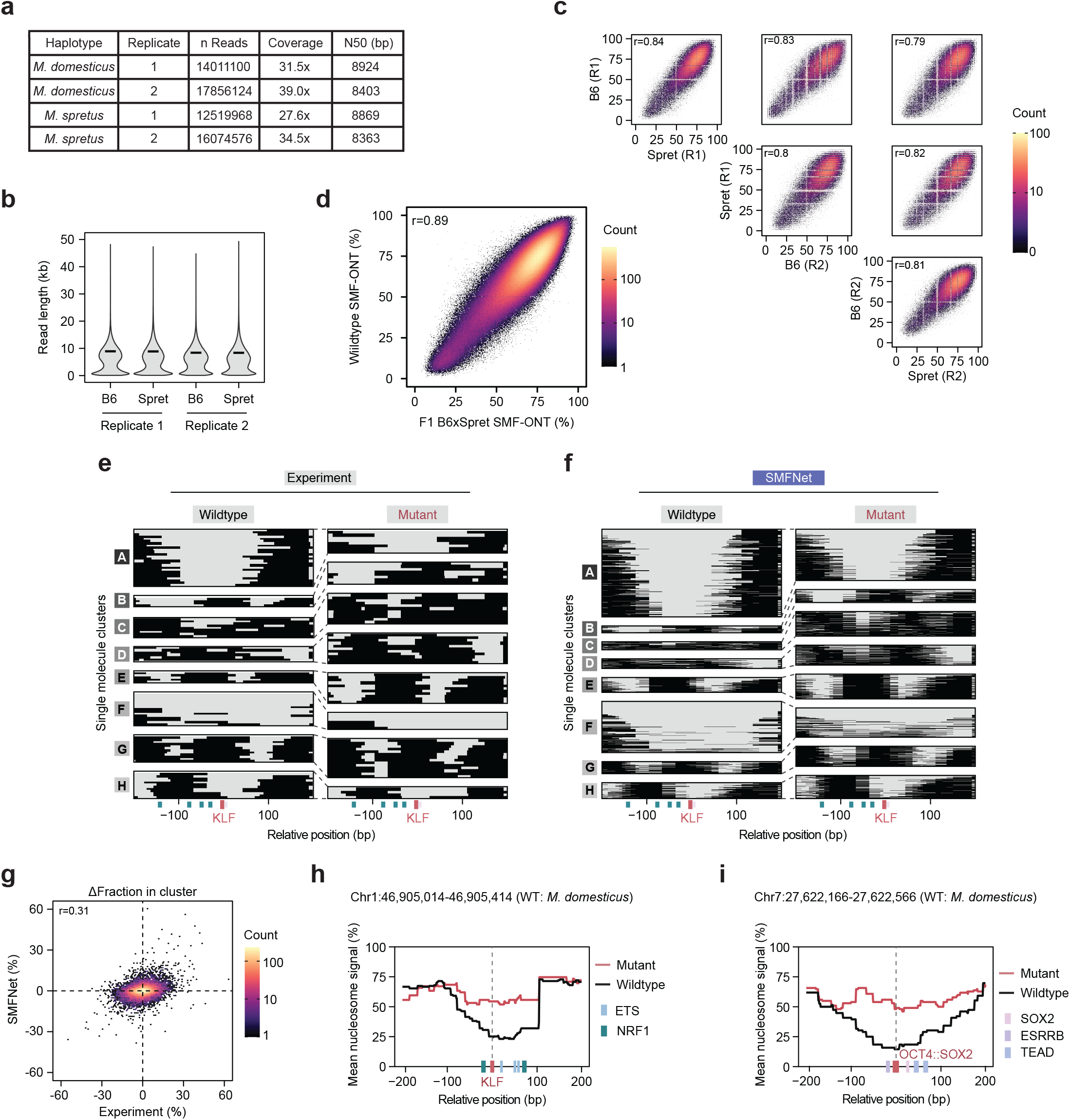
Allele-specific SMF-ONT and benchmarking of predicted variant effects. **a**) Summary statistics of long-read SMF sequencing data from F1 B6xSpret DNMT-TKO mESCs, shown separately by replicate and haplotype. Coverage was calculated as the cumulative length of filtered reads relative to the mouse genome size. **b**) Read-length distributions of long-read SMF data from TKO mESCs, shown by replicate and haplotype. Black bars indicate the N50 read length. The distributions were calculated from 100,000 randomly subsampled reads. **c**) Average SMF signal at 233,724 SMF positions across biological replicates and haplotypes (On chromosomes 1, 3 and 6). Only positions covered by at least 40 reads in all replicates and haplotypes were included. Pearson correlation coefficients are indicated. **d**) Average SMF signal across 1,762,359 positions in the F1 B6xSpret long-read SMF dataset compared to the wildtype long-read SMF data (On chromosomes 1, 3 and 6). Only positions covered by 100 reads in both datasets were included. The Pearson correlation coefficient is indicated. **e-f**) Experimental (**e**) and SMFNet-generated (**f**) single molecules at an endogenous enhancer (chr6:124,399,844–124,400,244), separated by wildtype (left) and mutant (right) allele. Molecules were clustered by nucleosome-occupancy profiles using Hamming distance. Accessible and nucleosome-occupied regions are shown in grey and black, respectively. SOX2 and KLF motifs are indicated in pink and teal, with the deleterious KLF variant in red. The wildtype allele is defined as the motif variant with the higher CWM score, indicating greater similarity to the optimal TF motif. **g)** Differences in single-molecule chromatin-state fractions between *M. domesticus* and *M. spretus* alleles across 2,212 regions centered on putatively deleterious TF motif variants (400 bp), comparing experimental measurements with SMFNet predictions. The Pearson correlation coefficient is indicated. **h-i**) Mean nucleosome profiles for *M. domesticus* and *M. spretus* alleles at representative loci disrupting a KLF (**h**, chr1:46,905,014-46,905,414) or OCT4::SOX2 motif (**i**, chr7:27,622,166-27,622,566). TF motifs are indicated by colored boxes below the plots, with the disrupted motif highlighted in red.

## Methods

### Experimental data generation

#### Cell culture

159 DNA methyltransferase triple-knockout (DNMT TKO) (100) and B6xSpret DNMT TKO (38, 101) mESCs were cultured on 0.2% gelatin-coated plates in ES medium (Dulbecco’s Modified Eagle Medium, supplemented with 15% FBS, 20 ng/ml leukemia inhibitory factor, 50 µM 2-mercaptoethanol, 2 mM L-glutamine and 1× nonessential amino acids) at 37 °C and 5% CO2. Medium was changed daily and cells were split every second day.

#### M.CviPI protein purification

The *M*.*CviPI* enzyme was produced as described previously (39). A detailed and updated purification protocol is supplied at protocols.io (102).

#### SMF-ONT

Single-molecule footprinting using Oxford Nanopore Technology (SMF-ONT) was performed as described previously (37, 39). Intact nuclei were extracted by re-suspending 2.5x10^5^ mESCs in ice-cold hypotonic lysis buffer (10 mM Tris pH = 7.4, 10 mM NaCl, 3 mM MgCl_2_, 0.1 mM EDTA, 0.5% IGEPAL). After 10 min incubation on ice, nuclei were spun down and washed with a wash buffer (10 mM Tris pH = 7.4, 10 mM NaCl, 3 mM MgCl_2_, 0.1 mM EDTA). Nuclei were spun down and resuspended in 1× M.GpC buffer (NEB, M0227L) using wide-bore tips. For treatment with *M*.*CviPI*, first reaction buffer was added (1× M.GpC buffer, 450 mM sucrose, 0.96 mM SAM (NEB, B9003S)) and the nuclei were then incubated with 200U of *M*.*CviPI* (EMBL PEPCORE), at 37 °C for 7.5 min. A second incubation round was performed with 100U of *M*.*CviPI* and 128 nmol of SAM (NEB, B9003S) at 37 °C for 7.5 min. A final incubation to methylate accessible CpG di-nucleotides was performed with 60U of *M*.*SssI* (NEB-M0226L), 3.5 µmol MgCl_2_ and 128 nmol of SAM at 37°C for 7.5 min. At each step, reaction tubes were mixed by inversion or flicking to avoid DNA shearing. Methyltransferases were heat-deactivated with a 20 min incubation at 65 °C and cooled down with 1 minute on ice. After an overnight Proteinase K (400 µg; 20 µL of a 20 mg/mL stock solution) digestion at 55 °C, high weight molecular DNA was extracted using the Quick-DNA HMW MagBead kit (Zymo, D6060) following the manufacturer protocol and using wide-bore tips. DNA concentration was measured using Qubit high sensitivity dsDNA kit (ThermoFisher, Q32854) and the DNA fragment size measured using FemtoPulse 165 kb kit (Agilent, FP-1002-0275) following manufacturer instructions. Ligation library kits (SQK-LSK114) were used with 2-2.5 µg of input DNA to generate the sequencing libraries. PromethION flow cells (R10.4.1) were loaded with 20-30 fmol of prepared library. Depending on the DNA fragment length and the DNA concentration, up to 3 reloads were performed per sequencing run using the flow cell wash kit (ONT, EXP-WSH004).

### Computational analysis

#### General data analysis and visualization

Unless otherwise stated, quantitative analyses were performed in *Python* v3.12 using *NumPy* v2.2.6 and *pandas* v3.0.5 (103, 104). Statistical summaries and data visualization were performed in *R* v4.4.1 using the *tidyverse* v2.0.0 packages (105).

#### SMF-ONT data processing

Nanopore data were processed using our previously described Nextflow pipeline, *nf-smfont* (39) (**Supplementary Table 1**). Reads were basecalled and aligned to the *Mus musculus* mm10 reference genome (*BSgenome*.*Mmusculus*.*UCSC*.*mm10*) using *Bonito* v0.7.3 with the super-accuracy model dna_r10.4.1_e8.2_400bps_sup@v4.2.0 and the integrated *Minimap2* v2.24 aligner. Reads with a quality score below 7 were discarded. Cytosine methylation was called using a published methylation model trained on CpG and GpC contexts (39). Methylation data was extracted from BAM files and converted into mBED format using pysam (33, 106). The resulting mBED files were indexed using *tabix* (HTSlib v1.19.2) (107). Methylation probabilities were transformed into log-likelihood ratios (LLRs), and cytosines with an absolute LLR of at least 2 were classified as methylated or unmethylated. Calls below this threshold were considered uncertain and excluded from downstream analyses. Reads with an overall methylation percentage below 5% or above 95% were also excluded. Before model training, methylation data was extracted from the mBED files in 2,114 or 2,048 bp windows and stored in a custom HDF5 format.

#### SMF-ONT data quality control

Read-length distributions and genome-wide coverage were assessed for each of the four biological replicates. Reproducibility between replicates was evaluated by calculating Pearson correlation coefficients for position-level methylation values. The SMF-ONT data were additionally compared with a short-read SMF dataset generated from the same cell type (22). Only genomic positions covered by at least 20 reads in each replicate were included in the correlation analyses.

### Model training and state inference

#### Model training and inference environment

SMFNet model training and sampling were performed with *PyTorch* v2.14.0 using *CUDA*-enabled GPUs (108). Computations were run on either a single NVIDIA H100 GPU or a single NVIDIA RTX 6000 GPU, depending on resource availability.

#### SMFNet model training

SMFNet predicts single-molecule profiles over the central 1,024 bp of a 2,048-bp DNA sequence (**Supplementary Note 2**). SMFNet comprises a 512-channel DNA encoder and a 128-channel denoising network, each with an initial kernel width of 21 and eight residual blocks with kernel width 9 and dilations 1–128. DNA embeddings conditioned every denoising block through feature-wise linear modulation (FiLM) (71). The denoising network accepts three-channel inputs encoding free and protected SMF states and a mask token, and uses a final single-channel convolution with kernel width 75. Residual blocks use layer normalization and ReLU. The jointly trained networks contain 21,189,505 parameters. Training used a masked discrete-diffusion objective to reconstruct masked SMF states at valid GpC/CpG positions, with a diffusiontime-weighted binary cross-entropy loss calculated against experimentally observed states.

Separate SMFNet models were trained using Illumina SMF (22, 69) and ONT-SMF data from 71,324 accessible and 132,518 inaccessible regions (**Supplementary Table 2**). Chromosomes 8 and 16 were used for validation, chromosomes 1, 3 and 6 for testing, and chromosomes 2, 4, 5, 7, 9–15, 17–19, X and Y for training. Optimization used AdamW with batch size 256, a peak learning rate of 10^−3^ with warm-up and cosine decay, and weight decay of 0.01, excluding biases. Training ran for a maximum of 400 epochs, with early stopping after 100 epochs without improvement in validation loss. The checkpoint with the lowest validation loss was selected.

#### SMFNet read sampling

Unless stated otherwise, we generated 1,024 synthetic SMF reads per region using a 32-step tau-leaping configuration, starting from fully masked reads (**Supplementary Note 3**). Remaining masked CpG/GpC reporter sites were selected uniformly for unmasking according to the diffusion schedule, with a minimum unmasking probability of 0.01. Newly unmasked states were drawn from Bernoulli distributions parameterized by the network’s sigmoid outputs.

#### ChromBPNet model training and motif annotation

To generate an ATAC-seq sequence-to-function model as a baseline for comparison with SMFNet, DNMT-KO mESC ATAC-seq reads were aligned as previously described using *Bowtie2* v2.5.4 (**Supplementary Table 2**) (99, 109). *ChromBPNet* v0.1.7 was trained using default parameters, with a peak set and data splits analogous to those used for SMFNet training (10). Base-pair-resolution attribution scores were computed using the DeepLIFT-SHAP implementation in *bpnetlite* v0.9.1 (64, 110), which was also used for model sampling. *TF-MoDISco* v0.5.16.0 was subsequently applied with default parameters to identify sequence motifs associated with chromatin accessibility (65). Finally, *Fi-NeMo* v0.30 was used to map the identified motifs to the genome, and the resulting motif instances served as annotations for all subsequent analyses (111).

#### SMF state calling

To identify TF footprints and nucleosome occupancy along individual experimental and generated SMF reads, contiguous methylated and unmethylated stretches were identified using custom Python code. Each stretch was extended halfway toward the nearest position with the opposite methylation state. Unmethylated stretches shorter than 70 bp were classified as bound, whereas stretches of at least 70 bp were classified as nucleosomal. Terminal stretches shorter than 50 bp were excluded to reduce edge effects.

### SMFNet benchmarking and dependency analysis

#### Quantification of SMFNet model performance

Agreement between observed and predicted single-molecule footprinting profiles was quantified across 13,716 ONT long-read and 11,814 Illumina short-read held-out test regions on chromosomes 1, 3, and 6 (**Supplementary Table 2, Supplementary Note 1**). Regions with at least 100 reads and 15 measurable cytosines were included. Mean signal agreement was assessed by Pearson correlation at individual positions and across 1,024-bp regions. For regional average profiles, we included a local sequence-context baseline using logistic regression trained on site-level mean SMF signals from training regions. The baseline predicted each site’s mean signal from the five flanking bases on either side, with separate models for CpG, GpC, and GCG sites. Agreement in positional dependencies was assessed by computing position-by-position correlation matrices and quantifying their similarity using Pearson correlation. Distributions of protected footprint lengths (consecutive unmethylated calls) were compared using the Wasserstein distance.

For each region, observed reads were randomly split once into two halves of equal size. Experimental pseudoreplicates compare one half with the other and give the agreement expected from a perfect model at that read depth. In the matched comparison, generated reads were subsampled to the size of one half and compared with the same observed half, so model and pseudoreplicate scores share read depth and reference. The full comparison uses all 1,024 generated reads against all observed reads. Position-shuffled experimental reads served as a control for protected run-length distributions.

Because substantially fewer positions were measurable per region in the short-read data, dependencies between positions were additionally quantified using pairwise log2 odds ratios. For the long-read model, frequencies of accessible, TF-bound, and nucleosomal state calls and mean SMF profiles were compared across 74,086 held-out putative TFBSs (**Supplementary Table 2**).

#### Co-accessibility analysis

State calls were collapsed by treating bound positions as accessible. Co-accessibility between position pairs was quantified as mutual information normalized by the geometric mean of their Shannon entropies, with the sign determined by Pearson correlation to distinguish concordant from discordant patterns. Pairs lacking variation or supported by fewer than 50 observed molecules were excluded. To correct for technical co-accessibility associated with molecule-to-molecule differences in overall methylation, the median score among the 20% most distant position pairs was subtracted as background. For visualization, position-wise scores were locally expanded and Gaussiansmoothed to generate continuous co-accessibility heatmaps. TFBS-level analyses were restricted to non-overlapping motif pairs separated by more than 15 bp, with each motif lying within 7.5 bp of a measurable cytosine.

For the composite CTCF analysis, 873 motifs in unseen test regions with at least 50 observed bound molecules were retained. Matrices were aligned to the nearest measurable SMF position and averaged across regions separately for observed and predicted bound and unbound states.

#### Co-binding analysis

TF co-binding was analyzed for non-overlapping motif pairs separated by >15 bp, with each TFBS located within 7.5 bp of an assayed cytosine (**Supplementary Table 2**). Pairs were required to have >5% TFBS occupancy and at least 100 observed molecules. Cobinding was quantified as the log2 odds ratio of the bound states. For single-molecule visualization, state calls were expanded to the midpoint between neighboring assayed positions.

#### Short-read GpC-only SMF models

SMFNet models were trained on short-read, single-enzyme SMF data from mouse embryonic stem cells, neural progenitor cells, C2C12 myoblasts, and MEL erythroleukemia cells (69). To distinguish accessibility labeling from endogenous CpG methylation, predictions were restricted to GpC sites without overlapping CpGs on either strand. Models used the standard SMFNet architecture and training procedure, with dropout of 0.2 and an exponential moving average of model weights with decay 0.999. Reverse-complement augmentation was omitted because the bait-capture data were strand-biased. Performance was evaluated on 28,360–30,693 held-out test regions using average-profile Pearson correlation and positional correlation matrix correlation as described above (**Supplementary Table 2**). For motif marginalization, selected TF motifs, derived from JASPAR annotations (112), were spiked into 200 background sequences with 50% GC content. Predicted signal was averaged across generated reads within each background, and the median signal across backgrounds was calculated at each position. Median profiles were smoothed using a rolling mean over 10 consecutive positions.

#### Fiber-seq model

SMFNet was trained on public Fiber-seq data from human GM12878 cells (35, 70) using the same architecture and training settings as the main model. Fiber-seq labels accessible DNA with m6A using *Hia5* methyltransferase and measures methylation on individual molecules by PacBio HiFi sequencing. The model predicted protection states at reference A and T positions, corresponding to adenines on opposite strands. Training regions comprised 2,114-bp windows centered on ENCODE cCREs, separated into regions overlapping GM12878 ATAC-seq peaks and background regions overlapping no peak (113). Predictions were evaluated in 17,384 held-out regions using Pearson correlations of average profiles and positional correlation matrices, as described above (**Supplementary Table 2**).

#### In silico mutagenesis

ISM was performed by independently substituting each nucleotide with all three alternative bases (**Supplementary Table 3**) (45). For ChromBPNet, the change in predicted total accessibility relative to the reference sequence was calculated for each substitution and averaged to obtain one score per genomic position. For SMFNet, a reference SMF profile was first predicted from a fully masked read conditioned on the reference DNA sequence. Each DNA position was then mutated to the three alternative bases, and the resulting changes in the predicted SMF profile were averaged across substitutions, yielding an influence matrix linking each mutated DNA position to every output position. Finally, the influence scores were summed across output positions to obtain a single positional ISM profile.

#### Benchmarking against ChromBPNet

Bulk SMFNet predictions were benchmarked against ChromBPNet (10) using background-marginalized motif insertions and predictions at endogenous TFBSs (**Supplementary Table 2**). For the insertion analysis, 20 chromatin-opening motifs identified by *TF-Modisco* v0.5.16.0 from the ChromBPNet model were inserted into 200 random background sequences (65). Accessibility was quantified as the percentage of non-nucleosomal SMFNet reads within ±25 bp of the insertion center and as the summed ChromBPNet signal within ±75 bp, respectively.

For endogenous regions, ISM scores were calculated across held-out test regions and evaluated at 169,683 putative TFBSs by summing the scores across all motif positions.

### Single-molecule chromatin state analysis

#### Clustering of single-molecule profiles

Clustering was performed on 400-bp windows centered on endogenous CREs from held-out test regions with at least 100 experimental reads, as well as on combinatorial motif perturbations at the *Phc3* promoter from a published mCHIRA dataset (**Supplementary Table 3**) (38). Sparse SMF state calls were converted to binary nucleosome-occupancy profiles and expanded to base-pair resolution. Experimental molecules with >15% missing values were excluded. Nearly fully protected molecules (>95% occupancy across 400 bp) were excluded as potential state-calling artifacts. This filter removed 2.66% of experimental and 3.12% of SMFNet-generated molecules at enhancer regions, and 2.48% and 0.92%, respectively, in the mCHIRA dataset. Equal numbers of experimental and SMFNet-generated molecules were jointly clustered using *HDBSCAN* v.0.8.44 with Hamming distance (114). Cluster medoids were defined as the molecule minimizing the summed within-cluster Hamming distance, and all molecules were subsequently assigned to their nearest medoid. Clustering was restricted to a maximum of 10 microstates for the endogenous CREs and 8 microstates for the mCHIRA analysis. Region-wide agreement between experimental and SMFNet-generated cluster-frequency distributions was quantified using Pearson correlation.

#### PRISM analysis

Per-read in silico mutagenesis (PRISM) was performed with SMFNet on individual single-molecule chromatin profiles. For each CRE, the corresponding 2,048-bp DNA sequence was provided together with the generated SMF read. For the mCHIRA analysis, positions outside the variable reporter sequence were masked. Each nucleotide within the target region was individually mutated to each of the three alternative bases, and the resulting change in model score was quantified using *single_read_ism* with one-position-at-a-time sampling. For each molecule, effects were averaged across eight diffusion trajectories (**Supplementary Note 6**). Differences in log probability between each substitution and the reference nucleotide were averaged across the three substitutions and visualized as sequence-logo attribution profiles of the wild-type sequence. For cluster-level analyses, PRISM attribution scores were averaged at each position across all molecules assigned to the corresponding chromatin state. TFBS-level PRISM scores were calculated as the mean attribution across the motif core and correlated with the corresponding change in cluster frequency following motif perturbation

#### Internal sampling assessment of PRISM predictions

To assess consistency between PRISM scores and model sampling, 128 genomic enhancer regions of 400 bp were randomly selected from held-out chromosomes (**Supplementary Table 3**). For each region, two experimental reads were randomly sampled and used as targets for PRISM. Following, the mutations with the most negative, most positive, and closest-tozero predicted effects were selected. Substitutions affecting CpG or GpC sites were excluded from the analysis. Predictions were validated by generating 1,024 SMFNet reads for each mutant and reference sequence. For each mutation, the observed effect was calculated as the difference in mean methylation agreement with the target read between mutantand reference-generated reads, considering only valid SMF positions in the target. Predicted and observed effects were compared using Spearman correlation. Additionally, mean changes in methylation agreement were calculated separately for the three mutation categories. Confidence intervals were estimated using 2,000 bootstrap replicates at the genomicregion level.

### *In silico* motif screening

#### ChIP-seq processing and GC-content analysis

Public mESC ChIP-seq datasets for KLF, MYC, OCT4, SOX2, ESRRB, REST, CTCF, ZIC3, NF-YA, NRF1, and BANP (**Supplementary Table 1**) (72, 86, 100, 115–119) were reprocessed as described previously (120) using *Bowtie2* v2.5.4 (121). Peaks were called using *MACS2* v2.2.7.1 (122). GC content was then calculated in 1-kb windows centered on ChIP-seq peak summits and summarized separately for each TF.

#### Synthetic sequence design and sampling

Consensus motifs of 20 non-redundant chromatin-opening TFs (**Extended Data Fig. E6a, Supplementary Table 4**) were inserted into 200 random background sequences using *tangermeme* v0.5.1 (123). Motifs derived from the ChromBPNet model that exceeded 16 bp were trimmed to the highest-scoring CWM window of that length. Except for the initial GC-content analysis (**Extended Data Fig. E6c**), backgrounds were generated at 50% GC content. Matches to any tested CWM motif in the background sequences were removed by local reshuffling. Motifs were tested individually, in heterotypic triplets comprising a central motif flanked by two identical partner motifs across variable spacings, and in homotypic arrays of 1–7 copies. Testing of heterotypic combinations was restricted to motif pairs occurring more than 1,000 times within 100 bp (motif-centre distance) in the SMFNet training data. For each sequence, 512 single-molecule chromatin profiles were generated with SMFNet. Predicted states were binarized into accessible and nucleosomal positions and expanded to base-pair resolution. Molecules with >95% occupancy of either state were excluded as potential state-calling artifacts, removing 3.12% of generated molecules across the in-silico libraries.

#### Quantification of accessibility

Molecules were considered accessible if no nucleosome was called within 7.5 bp of the sequence center. Accessibility was characterized by the fraction of accessible molecules, the median width of the central accessible region, and the precision of nucleosome phasing across molecules. Nucleosome phasing was quantified as boundary dispersion, defined as the median Euclidean distance between nucleosomal boundary positions across accessible molecules for a given sequence (**Supplementary Table 4**). For downstream analyses, all three measures were summarized as the median across backgrounds. For the GCcontent and motif-spacing analyses, accessibility frequency was additionally z-score normalized within each motif across the tested conditions by subtracting the mean and dividing by the standard deviation.

To visualize single-molecule distributions, profiles were pooled across backgrounds for each motif combination, ranked by overall nucleosome occupancy, and divided into 15 equally sized bins. Mean nucleosome occupancy was then calculated at each position within each bin.

#### Comparison with combined-effect baseline

SMFNet predictions for heterotypic motif triplets were compared with baselines derived from position-matched centre-only and pairedflank insertions (**Supplementary Table 4**). Treating paired flanks as one component incorporates their joint effect without isolating a pure three-way interaction. For each background and spacing, the combined construct BAB was compared with the central motif alone (−A−), paired flanks alone (B−B), and empty background (−−−). For a component X with accessible fraction p_X_ and background fraction p_0_, normalized accessibility gains and losses were u_X_ = max(p_X_ − p_0_, 0)/(1 − p_0_) and v_X_ = max(p_0_ − p_X_, 0)/p_0_, respectively. The component transformation was T_X_(p) = p(1 − v_X_) + (1 − p)u_X_. The compositional baseline averaged the outcomes of applying centre (A) and paired-flank (F) transformations in both orders: p_exp_ = {T_A_[T_F_(p_0_)] + T_F_[T_A_(p_0_)]}/2. These transformations combine predicted accessibility responses, rather than measured chromatin-transition rates.

When both components increase accessibility, the baseline simplifies to p_exp_ = p_A_ + p_F_ − p_0_ − (p_A_ − p_0_)(p_F_ − p_0_)/(1 − p_0_). The final term accounts for overlapping gains. Thus, accessibility below the uncorrected sum therefore need not imply antagonism. For positive-valued accessible-region width and boundary dispersion, effects were combined relative to background on the logarithmic scale: x_exp_ = x_A_ * x_F_/x_0_. For the frequency baseline, closing was set to zero at p_0_=0 and opening to zero at p_0_=1. Log-additive baselines required finite, positive control values. Invalid comparisons were excluded without pseudocounts. No such boundary cases or exclusions occurred in the analyzed data.

Expectations were calculated separately for each background and spacing. Joint predictions and expectations were summarized as medians across 200 backgrounds, and departures were calculated as differences between these medians, without mean-centering across motif combinations. Accessibility departures were expressed in percentage points. Agreement was assessed separately by metric and spacing using Pearson correlation and mean signed error, with confidence intervals estimated from 2,000 bootstrap resamples of backgrounds. These heuristic baselines identify departures from the specified compositional assumptions without establishing direct TF interactions.

#### Quantification and analysis of putative bound footprints

To assess how neighboring motifs influence TF binding, putative footprints were quantified at central OCT4::SOX2 motifs flanked by KLF or ESRRB motifs. These flanking motifs produce no measurable footprints above the background, reducing the risk of overlapping protection being assigned to the center motif. CG/GC sites within the inserted flank motifs were excluded from footprint calling. A putative bound footprint required protection at the single measured GC sites in the central OCT4::SOX2 motif and accessible sites on both sides separated by less than 70 bp. Total footprint frequency was calculated among all analyzed molecules, and conditional footprint frequency among the subset classified as centrally accessible. Each was counted directly. Conditional footprint frequency was not inferred by dividing the total frequency by accessibility. Changes upon adding flanks were measured relative to matched center-only constructs, and 95% confidence intervals were estimated using 2,000 bootstrap resamples of background sequences.

#### Comparison with OCT4 ChIP-seq predictions

*BPNet* v0.4.0 was trained on 17,192 mESC OCT4 ChIP-seq peaks using a published dataset and the bpnet-refactor workflow (23, 72). Training used default settings, except that the learning rate was set to 10^−4^, early-stopping patience to 10 epochs, and reverse-complement augmentation and training-data shuffling were enabled (https://github.com/kundajelab/bpnet-refactor). For the same synthetic sequences, we compared the change in BPNetpredicted OCT4 counts upon adding KLF or ESRRB flanks with the change in SMFNet total central-footprint frequency. Changes were summarized as medians across 200 backgrounds at each spacing, and agreement across spacings was assessed using Pearson correlation. This comparison tests concordance between model-predicted motif effects. It does not directly validate the synthetic constructs experimentally, and protection at OCT4::SOX2 motifs does not uniquely identify OCT4 occupancy.

#### Motif combination effects on nucleosome phasing

For each background and spacing, we calculated the changes in boundary dispersion and accessible fraction between combined and position-matched center-only constructs. These paired changes were summarized as medians across 200 backgrounds for each motif combination and spacing. We fitted a linear regression of median dispersion change against median accessibility change across all retained combinations and spacings. Regression residuals were reported as accessibility-adjusted dispersion changes. Negative residuals indicate lower dispersion, or more precise positioning, than expected from the fitted association with accessibility. This adjustment does not establish mechanistic independence from accessibility. Confidence intervals were estimated using 2,000 bootstrap resamples of background sequences.

### Allele-specific SMF analysis

#### F1 SMF-ONT data processing

SMF-ONT reads generated from B6xSpret DNMT TKO mESCs were processed as described above and aligned to the *Mus musculus* mm10 reference genome using *nf-smfont* (**Supplementary Table 1**) (39). Reads with an overall methylation level below 7.5% or above 95% were excluded. Known SPRET/EiJ SNP positions from the Mouse Genomes Project (124) were used as candidate variants for haplotype assignment. For each replicate, genotypes at these positions were called from the aligned ONT reads using *BCFtools* v1.21 (107). Heterozygous variants were then phased chromosome-wise with *WhatsHap* v2.6 (125) using read-level linkage information, and the resulting phased VCFs concatenated and used to haplotag individual reads. Reads were split by HP and PS tags using *pysam* v0.23.1 (106, 107), with unassigned reads excluded. Haplotype-specific BAM files were then re-processed with *nf-smfont* (39) to generate indexed mBED files for downstream SMF analysis.

#### F1 SMF-ONT quality control

Read-length distributions and genome-wide coverage were assessed separately for each haplotype and replicate. Reproducibility was evaluated by Pearson correlation of position-level methylation values across 233,724 SMF positions on chromosomes 1, 3, and 6, requiring at least 40 reads in each replicate–haplotype combination. In addition, the combined F1 dataset was compared with the wild-type DNMT-TKO SMF-ONT data using 1,762,359 positions covered by more than 100 reads in both datasets.

#### Parental variant calling

Candidate SPRET variants were genotyped from haplotagged F1 SMF-ONT reads across ATAC peaks, including 1,024 bp flanking sequences (99). Known SPRET/EiJ variants from the Mouse Genomes Project (124) were assigned to parental alleles when supported by at least five reads and 80% allele-consistent reads. Only variants confidently genotyped in both B6 and Spret were retained. Ambiguous variants were excluded.

#### Allele-specific motif annotation

To identify TF motifs affected by genetic variation, allele-specific B6 and Spret sequences were reconstructed by introducing confidently genotyped variants into the mm10 reference, including indels.

ChromBPNet hypothetical contribution scores were calculated separately for each allele using *DeepLIFT-SHAP*, and *Fi-NeMo* v0.30 was run independently on B6 and Spret sequences (10, 64, 111). Motif hits were projected to mm10 coordinates, and the homologous sequence on the opposite allele extracted and scored with the same contribution weight matrix (CWM). CWM scores were calculated as the sum of nucleotide-specific contribution weights across the motif sequence relative to the *TF-MoDisco* CWMs used for *Fi-NeMo* hit calling (65, 111).

Corresponding B6 and Spret motif hits were matched based on motif identity and genomic position, and a single representative annotation was retained while preserving allele-specific sequence and CWM information (**Supplementary Table 5**). Overlapping motif annotations were resolved by preferentially retaining the highestconfidence *Fi-NeMo* hits.

#### Selection of variant-containing motifs

We restricted the analysis to SNVs and indels located within the central 1,024 bp of each ATAC region. For each TF motif, variants were ranked by the difference in CWM score between alleles, and the top 10% with the largest differences were retained (**Supplementary Table 5**). To avoid multiple variants from the same regulatory region, only the variant with the largest CWM-score difference per region was analyzed.

#### Quantification of allele-specific chromatin accessibility metrics

Accessibility frequency, accessible-region width and boundary dispersion were quantified as described for the *in silico* analyses. Briefly, molecules were classified as accessible using a 15-bp window centered on the motif. Molecules with >95% occupancy of either state were excluded. In order to reliably estimate median width and boundary dispersion, analyses were additionally restricted to motifs with at least 20 accessible molecules in both B6 and Spret. Median accessible-region width was determined from the contiguous accessible segment surrounding the motif. Boundary dispersion was calculated from the pairwise euclidean distances between the left and right boundaries of accessible regions across molecules. Allele-specific effects were calculated as the difference between the two parental alleles. For all analysis, the allele containing the motif with the higher CWM score was defined as the wildtype allele.

#### Variant-effect prediction

SMFNet was used as described above to generate 512 molecules for variant-containing regions on held-out test chromosomes using the corresponding B6 and Spret sequences (**Supplementary Table 5**). Regions were selected based on allele-specific CWM differences as described, and predicted molecules were projected to mm10 coordinates. Analyses were restricted to regions with >50 experimentally observed reads per allele. The fraction of accessible molecules was calculated as described. In addition, 400-bp windows centered on the variant-containing motif were clustered using Hamming distance and *HDB-SCAN* v.0.8.44, with a maximum of eight clusters per region (114). In detail, equal numbers of experimental and SMFNet-generated B6 and Spret molecules were jointly clustered, and all remaining molecules subsequently assigned to the resulting cluster medoids. To assess prediction of allele-specific chromatin-state changes, differences in cluster frequencies between B6 and Spret were compared between experimental and SMFNet-generated molecules.

## Supplementary Notes

### Supplementary Note 1: Why and how SMFNet predicts chromatin-state distributions

SMFNet is a generative model trained to predict the distribution of single-molecule footprinting (SMF) reads in a genomic region from its DNA sequence. Modeling the full distribution, rather than the average accessibility at each position, matters because many different read distributions share the same average. Consider an enhancer with binding sites for two transcription factors (TFs), each bound on half of the molecules. If the TFs bind cooperatively, half of the molecules carry both footprints and the rest carry neither. If they bind exclusively, every molecule carries exactly one. The average accessibility at each site is the same in both cases, so bulk measurements such as ATAC-seq cannot tell them apart. Sequence models trained on bulk data, such as ChromBPNet (1) and AlphaGenome (2), have the same limitation because they predict the average signal at each position, treating positions as independent given the DNA sequence. Single SMF reads do distinguish the two cases, and a model that generates reads can capture which footprints occur together on the same molecule.

SMFNet learns read distributions with masked discrete diffusion (3), which trains a model on a masked prediction task similar to masked language modeling. We call the sites an assay can label *SMF positions*: GpC and CpG cytosines for the long- and short-read SMF data, GpCs that do not overlap a CpG for the GpC-only data, and A and T bases for Fiber-seq. During training, a random fraction of the SMF positions in each observed read is masked, from almost none to all of them, and SMFNet learns to predict the methylation state of the masked positions from the DNA sequence and the unmasked positions of the same read. After SMFNet has been trained, it can generate reads for new DNA sequences. Starting from a fully masked read, it predicts and unmasks a few positions at a time, and each later prediction conditions on the positions already unmasked. Footprints generated early therefore shape those generated later, which is how a generated read can contain both footprints of a cooperative pair, or only one of an exclusive pair. Supplementary Note 2 describes the architecture and training, and Supplementary Note 3 the sampling procedure.

#### 1.1 Evaluation metrics

We summarize how well generated reads match observed reads in a genomic region with three metrics, each computed per test chromosome region and then summarized across regions by their mean or distribution.

- **Profile Pearson** *r*: the Pearson correlation between the mean generated and mean observed SMF profiles across the region’s SMF positions.
- **Correlation of correlations**: the Pearson correlation between the pairwise position–position correlations of the generated reads and those of the observed reads, which measures how well SMFNet captures co-occurrence of protection within reads.
- **Ones run length Wasserstein**: the 1-Wasserstein distance between the distributions of lengths, in bp, of maximal runs of consecutive protected (unmethylated) positions in generated and observed reads.

Because each region has a finite number of observed reads, even a perfect model would not reach a profile correlation of one or a Wasserstein distance of zero. To show the best achievable value, we compute a pseudoreplicate optimum by splitting each region’s observed reads into two disjoint halves, of at most 1,024 reads each (the number of generated reads), and scoring one half against the other. Model metrics are scored against the same observed half, with a model sample of the same number of reads, so the pseudoreplicate gives the expected agreement of a perfect model at that read count. A site contributes only if at least 10 reads cover it in each observed half, a correlation of correlations uses every pair of positions with calls at both positions in at least 10 reads of each sample, excluding pairs where either position does not vary across those reads, and a metric is reported only when at least 10 sites, pairs, or runs remain. Run lengths are measured from the first to the last call of a maximal run of protected calls. A missing call ends a run, and runs touching the edge of the window are dropped. Before run lengths and position-pair correlations are computed, each generated read takes the missing-call pattern of a randomly chosen observed read, so that the model is not credited for denser coverage.

### Supplementary Note 2: SMFNet model architecture

The model consists of two modules, a DNA encoder *f*_*ϕ*_ and an SMF predictor *g*_*θ*_ that predicts the masked positions of a read (Supplementary Fig. 1). They are separated for training and sampling efficiency, as the DNA encoder can be run once per region while the SMF predictor can be updated using several reads for that region during training (Supplementary Note 2.1). Similarly, during sampling the SMF predictor has to be re-run each sampling step whereas the DNA encoder only has to be run once.

Both modules are dilated convolutional neural networks (CNNs) inspired by the ChromBPNet architecture and combined have 21.2 million trainable parameters. Each module starts with an input convolution with kernel size 21, followed by eight dilation blocks with dilations *d* = 1, 2, 4, …, 128. Each dilation block applies layer normalization, a dilated convolution with kernel size 9, and a ReLU, and adds the result to the block’s input. In the SMF predictor, each block first conditions its input on the DNA embedding with FiLM (4), which scales and shifts each channel using a projection of the embedding.

The DNA encoder takes as input a one-hot encoding of the 2,048-bp DNA sequence, where bases are integer encoded with A, C, G, T = 0, 1, 2, 3 and *−*1 for unknown bases (N), which one-hot encode to all zeros. Its dilation blocks have 512 filters and produce a 2,048 by 512 dimensional embedding, which is cropped to the central 1,024 bp and conditions every dilation block of the SMF predictor.

The SMF predictor takes as input a partially masked read over the central 1,024 bp, one-hot encoded so that zero corresponds to methylated (accessible), one to not methylated (protected), and two to masked or missing, giving a 1,024 by 3 dimensional input. Its dilation blocks have 128 filters, and an output convolution with kernel size 75 produces 1,024 logits, which are trained to predict the values of the masked positions of the input read.

During training, we sample 32 reads for each DNA sequence. As the DNA sequence is the same for those reads, we only run the DNA encoder once and broadcast its embeddings to the SMF predictor. This same technique is used during sampling, enabling us to reuse the DNA embeddings across sampling steps and multiple reads for the same DNA sequence.24 x 512)

**Supplementary Fig. 1.**
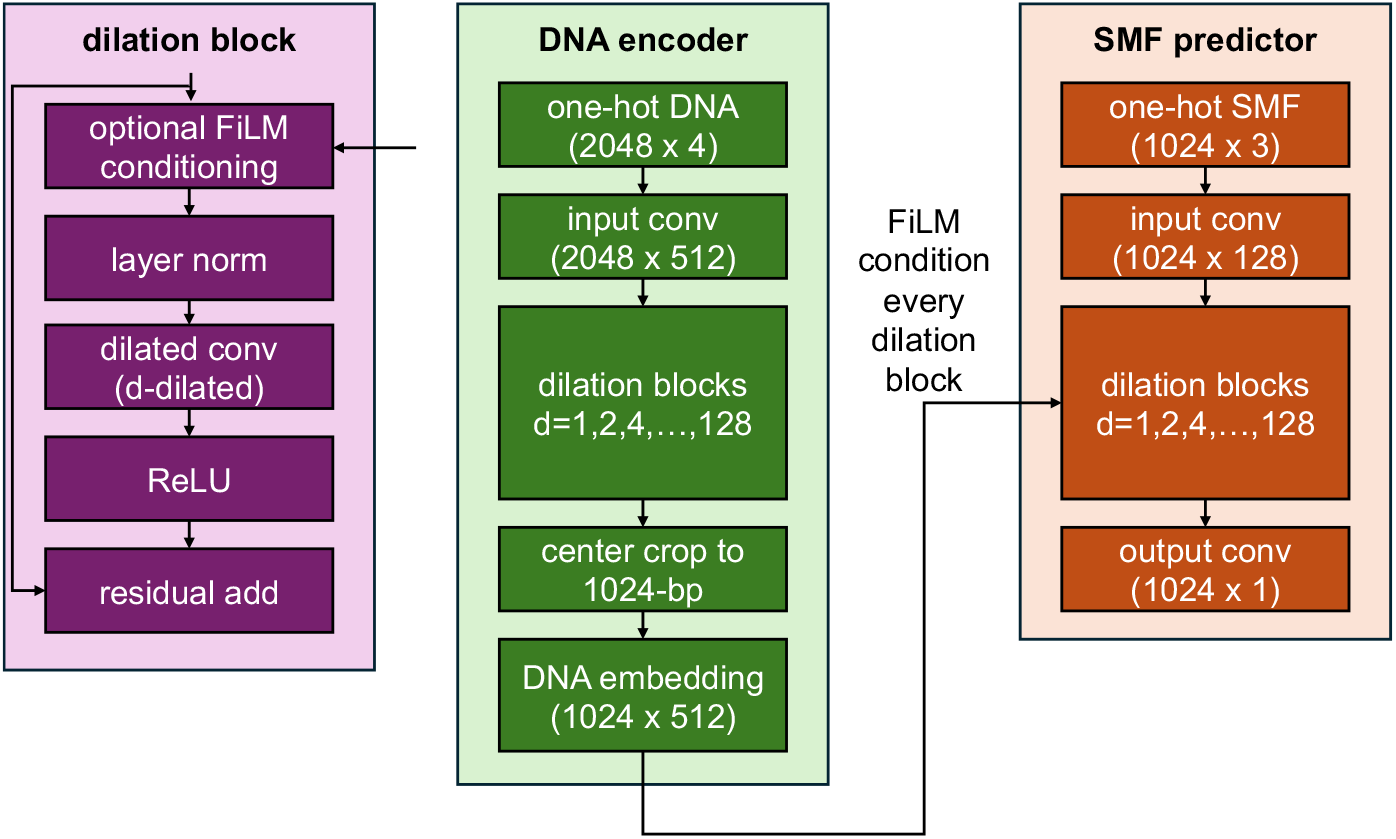
SMFNet architecture. The DNA encoder *f*_*ϕ*_ (middle) embeds the one-hot encoded 2,048-bp DNA sequence, and the central 1,024 bp of the embedding condition every dilation block of the SMF predictor *g*_*θ*_ (right) through FiLM. The SMF predictor takes a partially masked read, with one channel each for methylated, not methylated, and masked positions, and outputs one logit per position for the probability that it is not methylated (protected). Left, a single dilation block, where FiLM conditioning is used only in the SMF predictor. Shapes are given as length *×* channels, and *d* is the dilation. The DNA embedding is computed once per region and reused across reads and sampling steps.

#### 2.1. Training SMFNet

SMFNet is trained for up to 400 epochs, with early stopping on validation loss, where each epoch corresponds to seeing each train set genomic region once. In each epoch a new set of 32 reads is selected randomly without replacement when the region has at least 32 reads and with replacement otherwise. Per epoch, the DNA encoder is trained on every available genomic region while the SMF predictor is trained on roughly 32 SMF reads per region. We summarize other training hyperparameters in Supplementary Table 1.

During training we also randomly reverse complement the data and select a randomly shifted window within each region as augmentations. Regions are stored as 2,114-bp windows, so each training example is a 2,048-bp crop at a uniformly random offset of 0 to 66 bp, while evaluation uses the centered crop. The GpC-only models use 2,048-bp bait windows, so they have no shift augmentation. Algorithm 1 shows pseudocode for a full training epoch.

**Supplementary Table 1.**
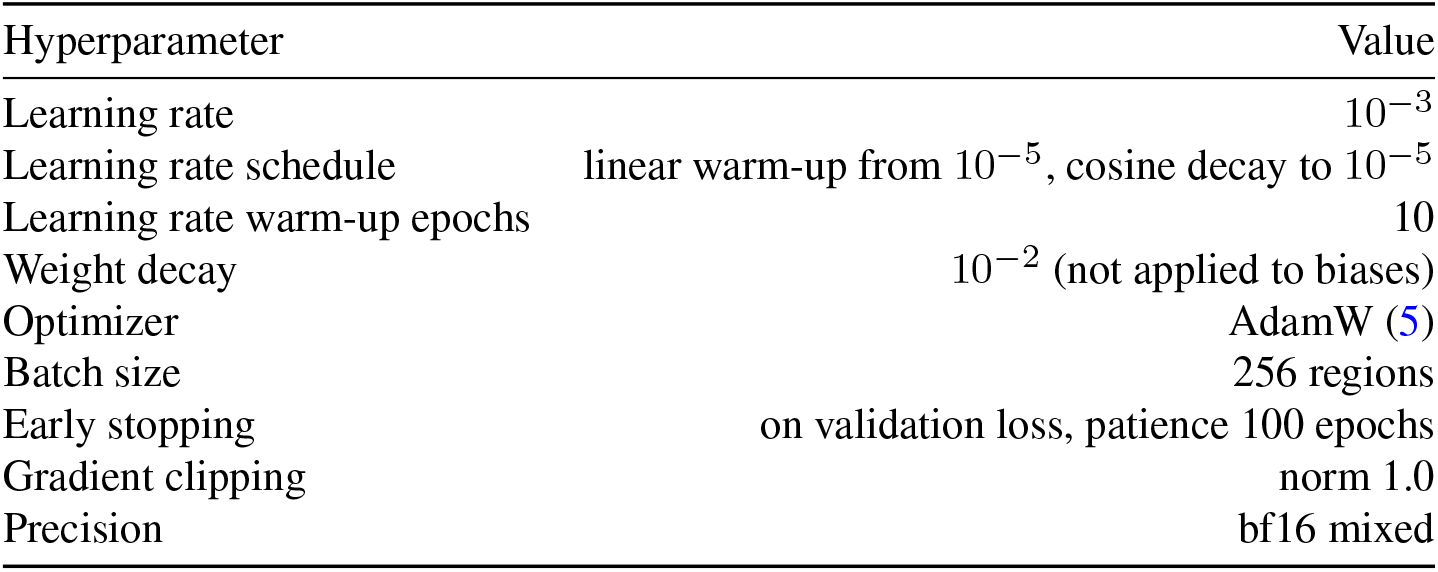
SMFNet training details.

| Hyperparameter | Value |
|---|---|
| Learning rate | 10<sup>−3</sup> |
| Learning rate schedule | linear warm-up from 10<sup>−5</sup>, cosine decay to 10<sup>−5</sup> |
| Learning rate warm-up epochs | 10 |
| Weight decay | 10<sup>−2</sup> (not applied to biases) |
| Optimizer | AdamW (5) |
| Batch size | 256 regions |
| Early stopping | on validation loss, patience 100 epochs |
| Gradient clipping | norm 1.0 |
| Precision | bf16 mixed |

**Supplementary Fig. 2.**
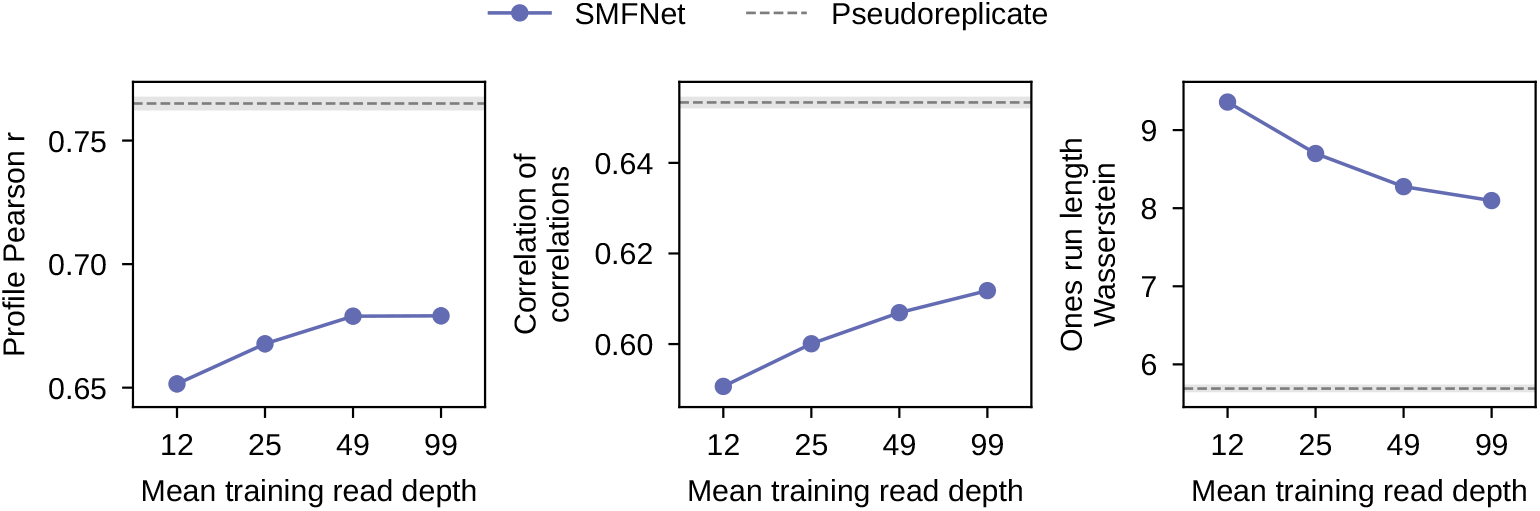
Effect of simulated reduced read depth on test set metrics, computed on all test regions. The x axis gives the mean training read depth per site after Bernoulli thinning to 1/8, 1/4, 1/2, and all reads (99 is the full model). While all metrics improve with greater read depth, the overall difference in performance is small even at 1/4 read depth (25 reads per site). Points show the mean over test regions and error bars the 95% bootstrap confidence interval of the mean (5,000 resamples of regions). Dashed lines and gray bands show the mean and 95% bootstrap confidence interval of the pseudoreplicate optimum, which uses only observed reads and so is the same for every setting.

#### 2.2. Effect of reduced read depth

We tested SMFNet’s performance when trained on simulated read depths of 1/8, 1/4, and 1/2 the number of train and validation chromosome reads used to train the full model (Supplementary Fig. 2). To simulate a given read depth fraction *β*, we used Bernoulli thinning in which each read was independently randomly retained with probability *β*. Thinning was applied once with a fixed random seed and only to the training and validation chromosomes, so the retained reads are nested across fractions and the test set is unchanged. Regions left without reads were dropped. Thinning to 1/8, 1/4, 1/2, and all reads gives a mean training read depth of 12, 25, 49, and 99 reads per site (reads with a call at a site, averaged over every site in the training split). All reported metrics were measured on the test chromosomes.

##### Algorithm 1

SMFNet training epoch

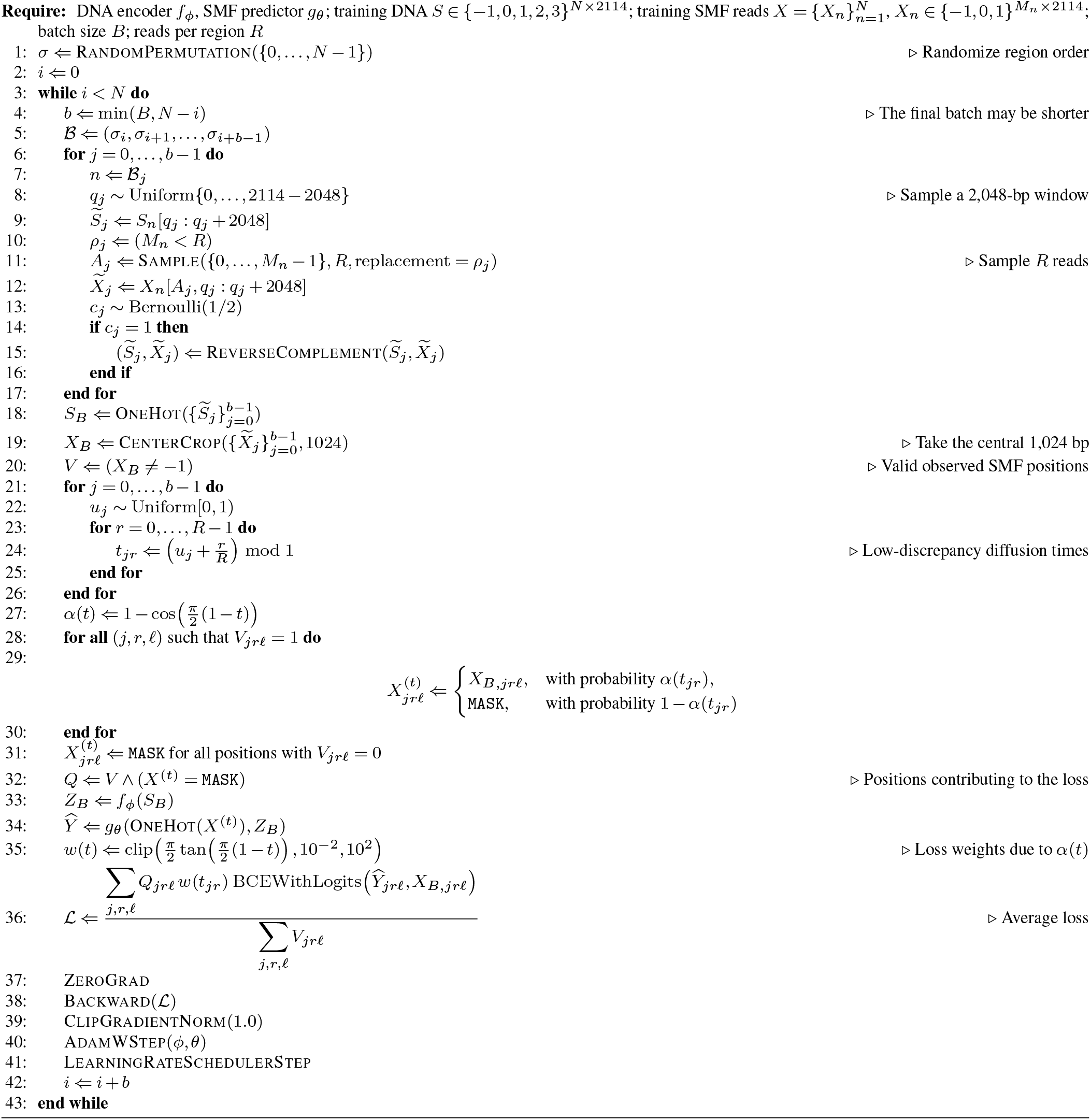

### Supplementary Note 3: Sampling from SMFNet

While the most accurate way to sample from SMFNet is to unmask only one SMF position at a time, this is unnecessarily computationally expensive as it requires a model forward pass for each SMF position. Instead, we use a standard sampling procedure for discrete diffusion models known as tau leaping (3). In tau leaping, one pre-specifies the number of sampling steps *T*, e.g. *T* = 32, and at each sampling step several SMF positions may be unmasked at once. The number of positions unmasked at each sampling step is set by an unmasking schedule. SMFNet uses the cosine unmasking schedule (3), defined by the function *α*(*t*) giving the expected fraction of SMF positions that are unmasked at diffusion time *t*,

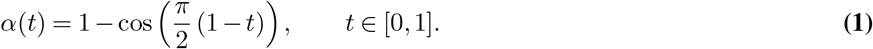

At *t* = 1 every position is masked (*α*(1) = 0) and at *t* = 0 every position is unmasked (*α*() = 1). At each sampling step *k* = *T*,…, 1 the probability of unmasking a still masked position is

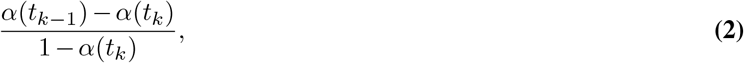

where *t*_*k*_ = *k/T* . The cosine schedule starts unmasking positions gradually at early sampling steps then unmasks more positions in parallel later in the sampling process when more context is available.

As SMFNet is only trained on SMF positions, we only unmask at those positions. If at sampling step *k* site *j∈{* 0, …, 1023} is randomly selected for unmasking according to Equation 2, its value is sampled according to SMFNet’s predicted probability for that site,

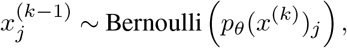

where *p*_*θ*_(*x*^(*k*)^)_*j*_ is SMFNet’s predicted probability that the *j*th site is unmethylated (protected) given *x*^(*k*)^. We provide pseudocode for the sampling procedure in Algorithm 2. In practice, the unmasking probability is floored at *ε* = 0.01, at least one masked position is revealed in each read at every step, and the final step reveals all remaining positions. All generated reads in this study use this sampler.

#### 3.1. Effect of number of sampling steps *T*

We evaluated the effect of the number of tau-leaping sampling steps *T* on test set metrics by generating reads with *T ∈* {16, 32, 64, 96, 128} (Supplementary Fig. 3). The profile Pearson correlation does not depend on *T*, whereas the correlation of correlations and ones run length Wasserstein distance improve with more steps up to *T* = 64 and change little beyond it. This is consistent with marginal accessibility being learned per position while within-read structure depends on how many positions are unmasked in parallel at each sampling step. We therefore use *T* = 32 by default, which captures most of the improvement in within-read structure at half the cost of *T* = 64 (Supplementary Fig. 3).

**Supplementary Fig. 3.**
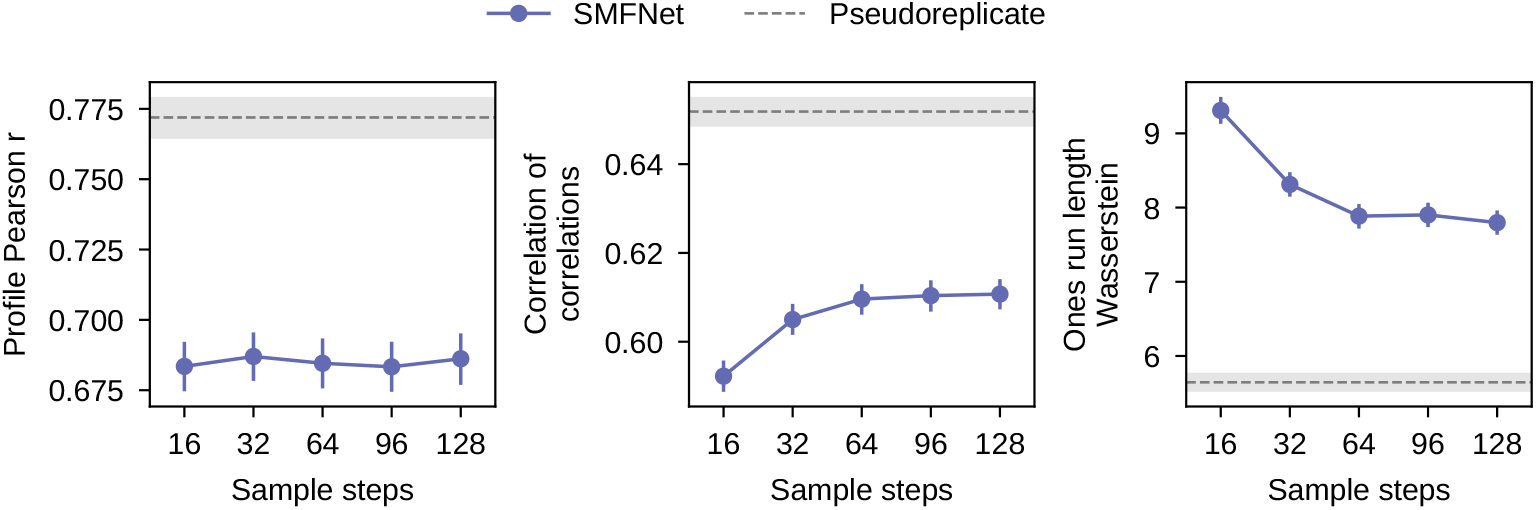
Effect of the number of tau-leaping sampling steps *T* on test set metrics, computed on a subset of 2,048 test regions. Points show the mean over test regions and error bars the 95% bootstrap confidence interval of the mean (5,000 resamples of regions). Dashed lines and gray bands show the mean and 95% bootstrap confidence interval of the pseudoreplicate optimum, which uses only observed reads and so is the same for every setting.

##### Algorithm 2

Tau-leaping sampling from SMFNet

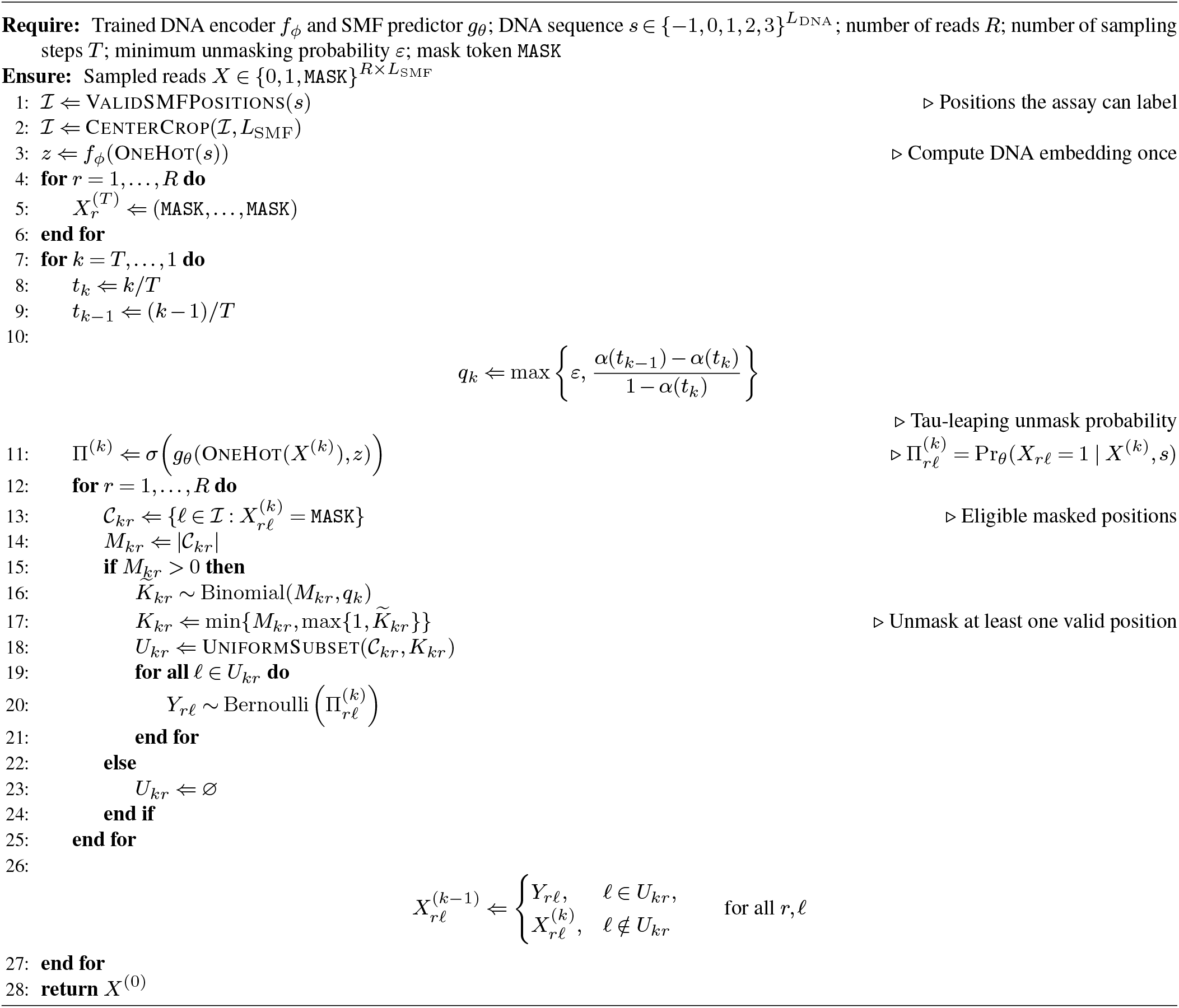

### Supplementary Note 4: Cell-type-specific accessibility around TF motifs

To test whether the cell line models (Methods, short-read GpC-only SMF models) capture differences in accessibility between cell types, we aggregated SMF signal around strong motif matches in the output windows of test regions (Supplementary Fig. 4). For each match to a JASPAR position weight matrix (PWM) (6) with a relative score of at least 0.90 (the log-odds score scaled between the matrix’s minimum and maximum, scanned on both strands), every eligible GpC within 100 bp of the motif center was placed at its strand-oriented distance from the center. Eligible GpCs are shared by all four cell types, with at least 30 observed calls across a region’s reads (21 in the evaluation subset described below) and at least 10 calls among the generated reads, and a region needs at least 10 such sites. For each cell type, we took the SMF signal at these sites (the percentage of protected reads) relative to that cell type’s average over the same region, so that region-wide differences between cell types cancel, and averaged it in 10-bp bins. The region average is the cell type’s mean SMF signal over all eligible GpCs in the region’s output window. Every pair of motif match and GpC counts equally, including overlapping matches, and confidence intervals resample whole regions so that all sites from a region stay together. We computed this separately for observed and generated reads. The observed signal uses a random evaluation subset of 70% of each region’s test set reads, and the generated signal uses reads from each cell type’s model. CTCF motifs show a similar footprint in all four cell types, whereas OCT4::SOX2 motifs are less protected, i.e. more accessible, in ES cells and GATA1::TAL1 motifs in MEL cells, in both observed and generated reads.

**Supplementary Fig. 4.**
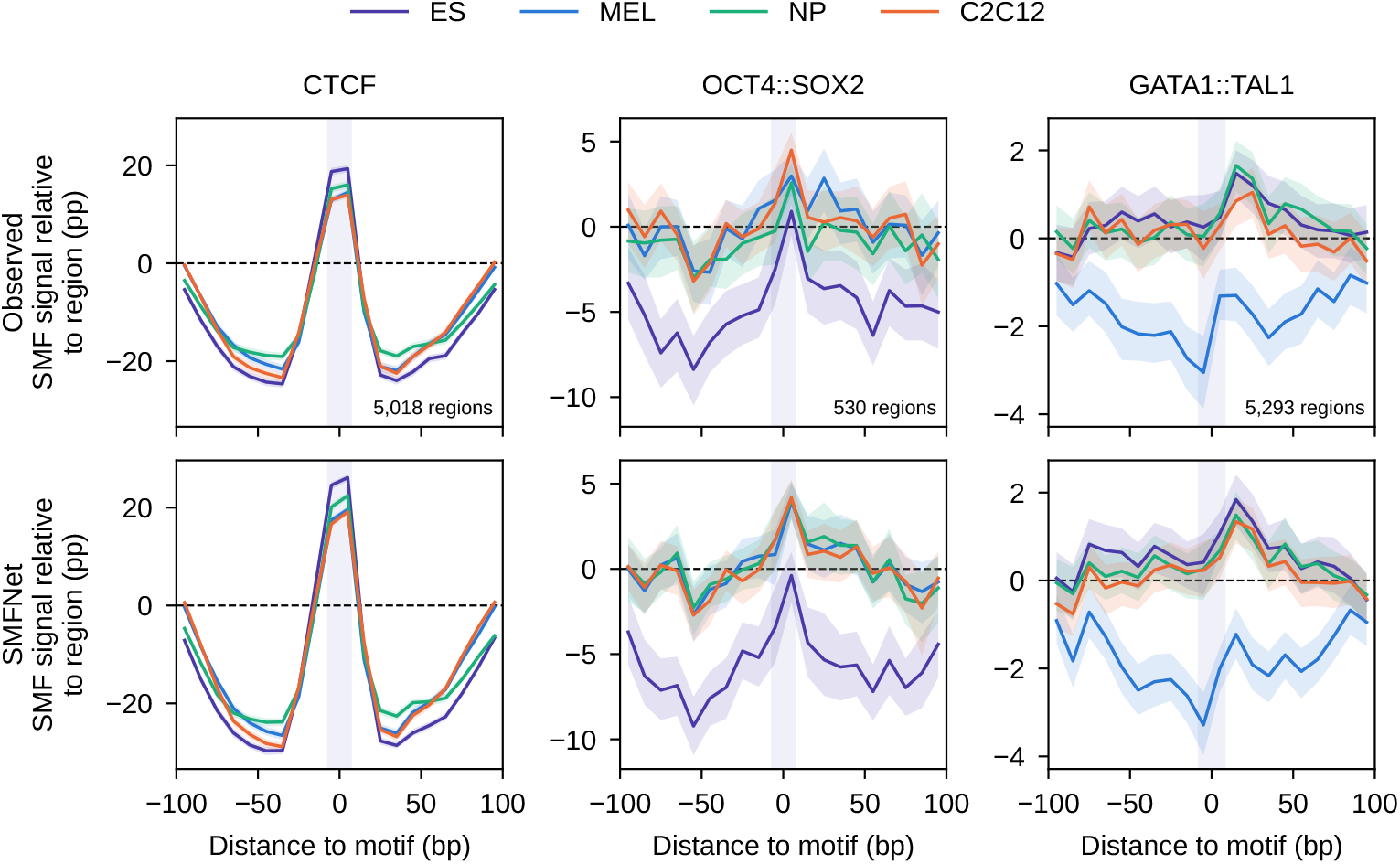
SMF signal around strong motif matches in test regions for each cell type, relative to the cell type’s average over the region, for observed (top) and SMFNet-generated (bottom) reads. Columns are motifs, JASPAR matrices MA0139.2 (CTCF), MA0142.1 (OCT4::SOX2), and MA0140.3 (GATA1::TAL1), each with its own y axis; the shaded band marks the motif. Lines are means over sites in 10-bp bins, and bands are 95% confidence intervals from 500 bootstrap resamples of regions. The number of regions with motif matches is given in each top panel. pp, percentage points.

### Supplementary Note 5: Fiber-seq example regions

To complement Extended Data Fig. E3, Supplementary Fig. 5 shows observed and SMFNet-generated Fiber-seq reads for three randomly selected test set regions. The Fiber-seq model and its training data are described in the Methods.

### Supplementary Note 6: Per-Read In Silico Mutagenesis (PRISM)

In silico saturation mutagenesis (ISM) interprets genomic deep learning models by substituting each base of a DNA sequence with every alternative base and recording how the model’s prediction changes (7, 8). ISM is usually applied to models such as ChromBPNet and AlphaGenome, which make a deterministic prediction from a sequence. SMFNet instead generates random reads (Supplementary Note 3), so ISM applied to SMFNet needs a quantity that depends deterministically on the sequence. Per-Read In Silico Mutagenesis (PRISM) estimates the effect of mutating each nucleotide on the log likelihood of generating a given SMF read.

SMFNet generates a read through a trajectory of unmasking steps that starts from a fully masked read and ends at a fully unmasked one. Let *x*^(*t*)^ *∈ {−*1, 0, 1*}*^1024^ be a partially masked read where, as in Supplementary Note 3, the sampling process runs backwards from *t* = *T* to *t* = 0, so *x*^(*T*)^ is fully masked and *x*^(0)^ is the fully unmasked observed read. Note that for any given read, we can construct many possible sampling paths *x*^(1)^, …, *x*^(*T−*1)^ depending on which positions get unmasked at each sampling step. PRISM addresses this by averaging PRISM scores over several sampling paths.

**Supplementary Fig. 5.**
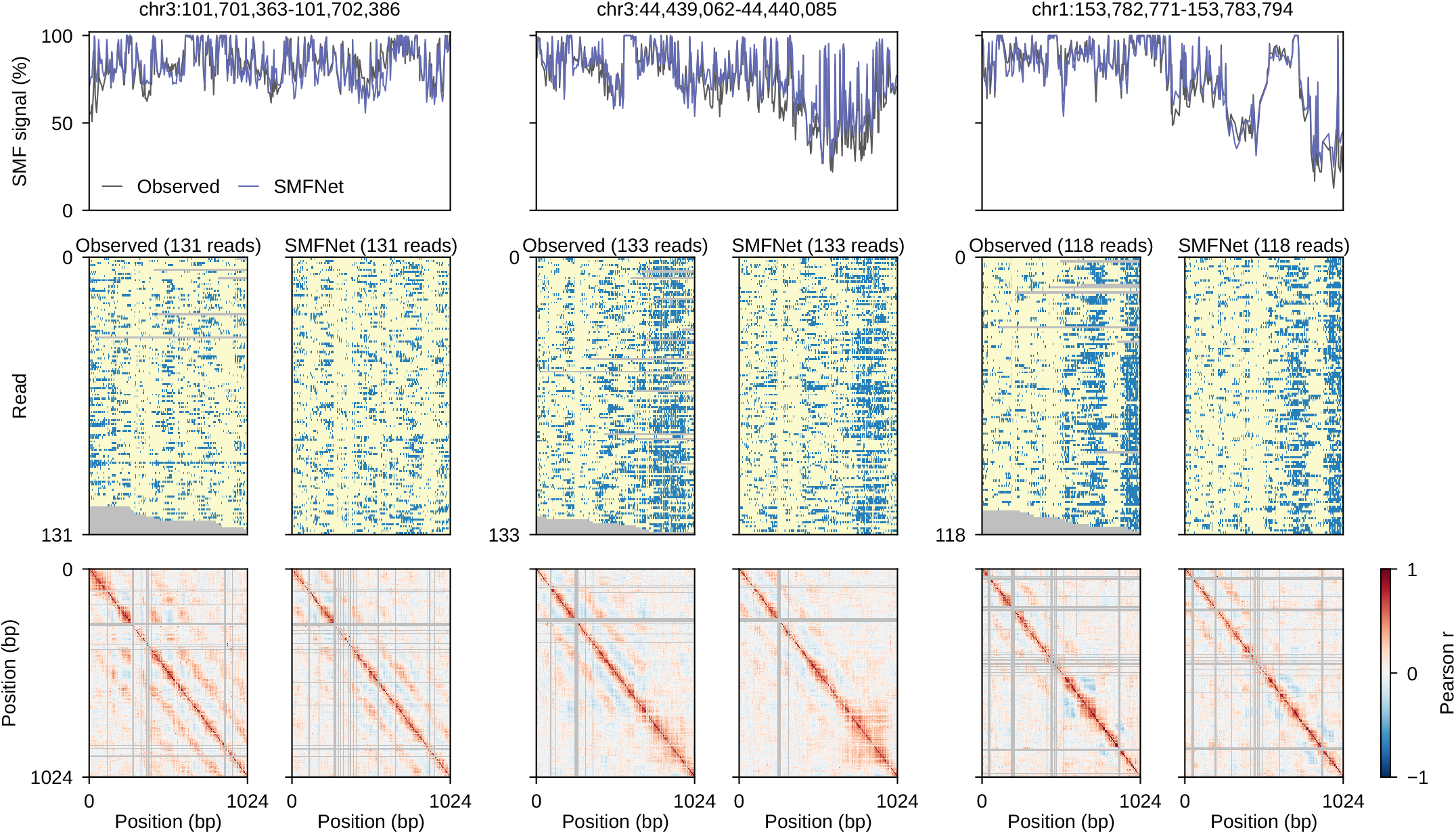
Observed and SMFNet-generated Fiber-seq reads in three randomly selected test set regions from human GM12878 cells (hg38). **Top** SMF signal (percentage of reads without m6A, i.e. protected) of observed (gray) and generated (purple) reads; the generated profile averages all generated reads for the region. **Middle** Observed reads and an equal number of randomly chosen generated reads. Blue indicates m6A (free) and yellow no m6A (occupied); gray marks missing calls. Each site spans to the midpoints with its neighbors, so all rows share the base-pair axis. **Bottom** Pairwise Pearson correlations between sites across the reads shown above, so observed and generated correlations use the same number of reads. Gray marks site pairs with fewer than 10 joint reads or without variation.

We define ℐ *⊂* {0, …, 1023} to be the set of valid read positions, i.e. *x*_*l*_ *∈* {0, 1} for all *l ∈ ℐ*, where *x* = *x*^(0)^ is the observed read. At each sampling step, SMFNet predicts the probability that each still-masked position is methylated or not methylated using the DNA sequence *s ∈ {−*1, 0, 1, 2, 3*}*^2048^. Given the set *U*_*t*_ of positions revealed when moving from step *t* to *t −* 1, the log probability that they take their observed values is 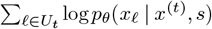. Summing over steps gives the log probability of the read along the trajectory *U* = (*U*_*T*_, …, *U*_1_),

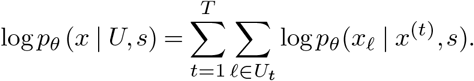

We construct trajectories in one of two ways, both starting from a random permutation *π* of the read positions. With *tau-leaping* trajectories, unmasking follows the cosine schedule of Supplementary Note 3 with *T* steps, so that *x*^(*t*)^ reveals the valid positions among the first ⌊*α*(*t/T*) *L*_SMF_⌋ entries of *π* (Algorithm 3). When positions outside a window are kept masked throughout, as in the PRISM validation (Supplementary Note 6.1), *π* ranks only the valid positions inside the window. At each step, these trajectories reveal the positions with the lowest ranks in a fixed unmasking order, as set by the cosine schedule, rather than random reveal counts as in Algorithm 2, and PRISM scores refer to these fixed trajectories. With *one-position-at-a-time* trajectories, each step reveals the next valid position in *π*, so *T* =|*ℐ*| and the trajectory score is the read’s log likelihood under the autoregressive factorization given by *π*. The PRISM validation uses tau-leaping trajectories with *T* = 32, whereas the analyses of individual chromatin states in the main text (Fig. 2) use one-position-at-a-time trajectories, which require more SMF predictor evaluations but match the most accurate sampling procedure (Supplementary Note 3).

Using these trajectory log probabilities, we can get the PRISM scores for each base substitution. Let *s*_*i→j*_ for *j ∈* {0, 1, 2, 3} denote the DNA sequence where every nucleotide stays as its reference value except position *i*, which is changed to base *j*. For a single unmasking order, we get a PRISM score matrix with one row per mutated position and one column per base,

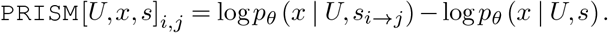

Note that when *s*_*i*_ = *j*, the PRISM score is zero. Unless stated otherwise, PRISM scores all three alternative bases at every mutated position. The validation (Supplementary Note 6.1) uses only substitutions that preserve the SMF position mask. This construction makes the PRISM score equal to the log-probability ratio of the read along trajectory *U* according to the model for the mutated sequence vs. the original sequence. In other words, the PRISM score is positive when the mutation makes sampling the given read with unmasking trajectory *U* more likely and negative if the mutation reduces the read’s likelihood. We provide pseudocode for PRISM in Algorithm 3 and a visual overview in Extended Data Fig. E4f.

The unmasking trajectory *U* has little effect on the score (Supplementary Fig. 6). For the mutations selected in the PRISM validation (Supplementary Note 6.1), scores under individual unmasking orders closely track their mean over orders, with a median standard deviation across orders of 0.09 and 99.5% sign agreement with the mean. For an example read, scores for every substitution are also highly correlated between orders (Pearson *r* of 0.71–0.90). We nonetheless average over *K* = 8 unmasking orders to get our final PRISM score 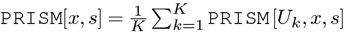.

A central limitation of PRISM is its computational cost. Scoring all substitutions at the central *L* positions of a region requires 3*L* + 1 passes of the DNA encoder, one for each substituted sequence and one for the reference, since each sequence’s DNA embedding is computed once and shared across all *K* unmasking orders. It also requires (3*L* + 1) *× K× T* evaluations of the SMF predictor, one per sequence, unmasking order, and sampling step. Because the unmasking trajectory is fixed by the observed read, the *T* evaluations for each sequence and order do not depend on one another and run as a single batch. With one-position-at-a-time trajectories, *T* is the number of valid read positions, so the SMF predictor dominates the cost. While PRISM can be easily batched on GPU and SMFNet itself is efficient, the computational cost currently makes PRISM better suited to analyzing particular regions than to genome-wide application.

#### 6.1. Quantitative PRISM validation

The Methods describe the PRISM validation (Extended Data Fig. E4h–i). Here we give the details of window and mutation selection, confirmation scoring, and the empirical mutation effect.

Each of the 128 enhancer regions was evaluated in a 400-bp window centered on a candidate enhancer motif cluster and fully contained in the model output, keeping the most central eligible window per source region. Windows required at least 25 observed reads with valid methylation calls at 25 or more positions and at least 90% of the SMF positions in the window. We scored only substitutions that preserve the SMF position mask across the model output. For each target read, mutations were selected using PRISM scores averaged over eight random unmasking orders, and the selected mutations were then rescored with eight independent orders without changing the selection. These confirmation scores are the ones reported. During both PRISM scoring and generation, SMF positions outside the window stayed masked while the full 2,048-bp DNA sequence was kept as context, and generation used 32 sampling steps. Reads generated from the same conditioning sequence were shared across the targets in a region.

##### Algorithm 3

Per-Read In Silico Mutagenesis (PRISM)

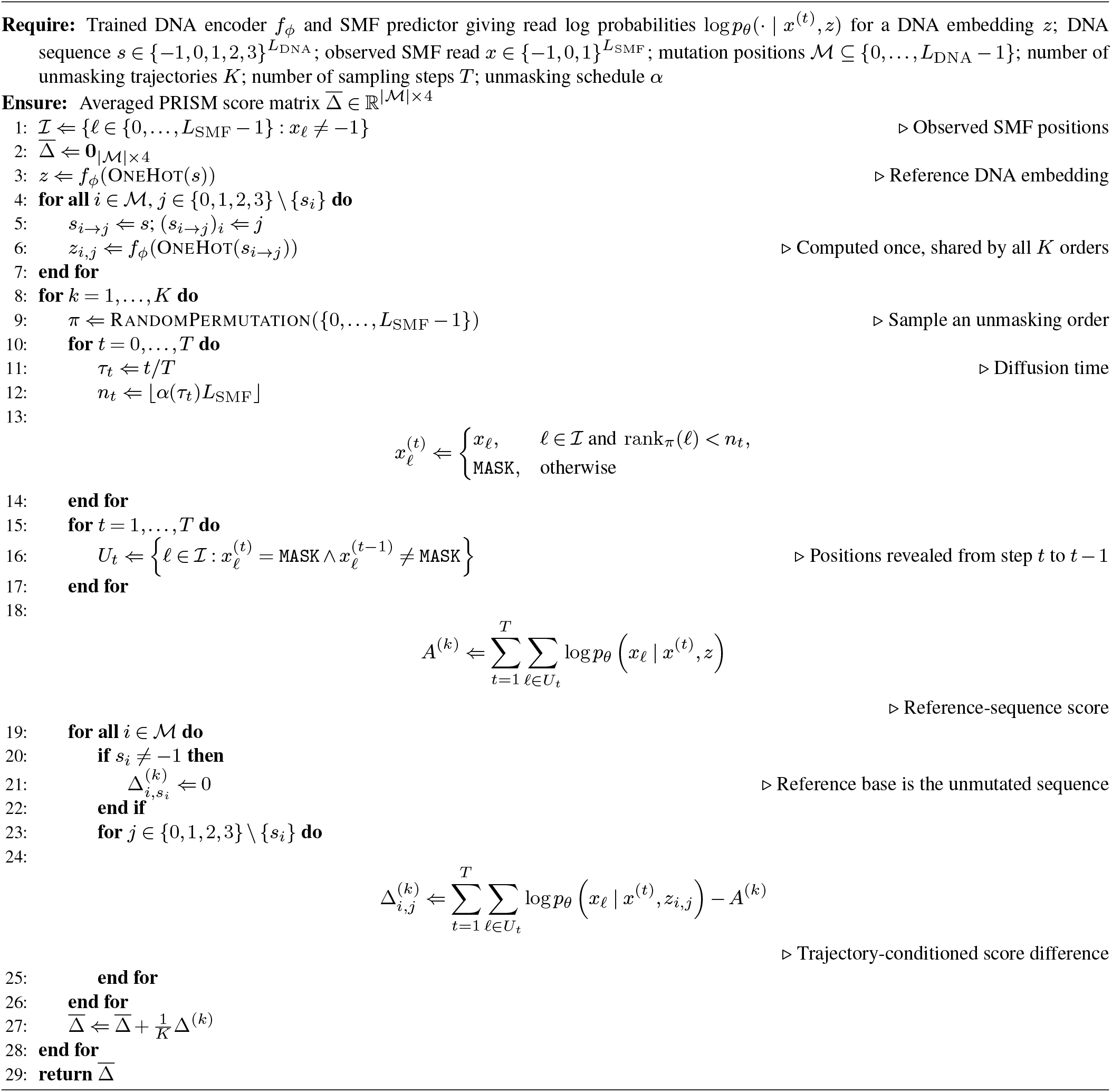

**Supplementary Fig. 6.**
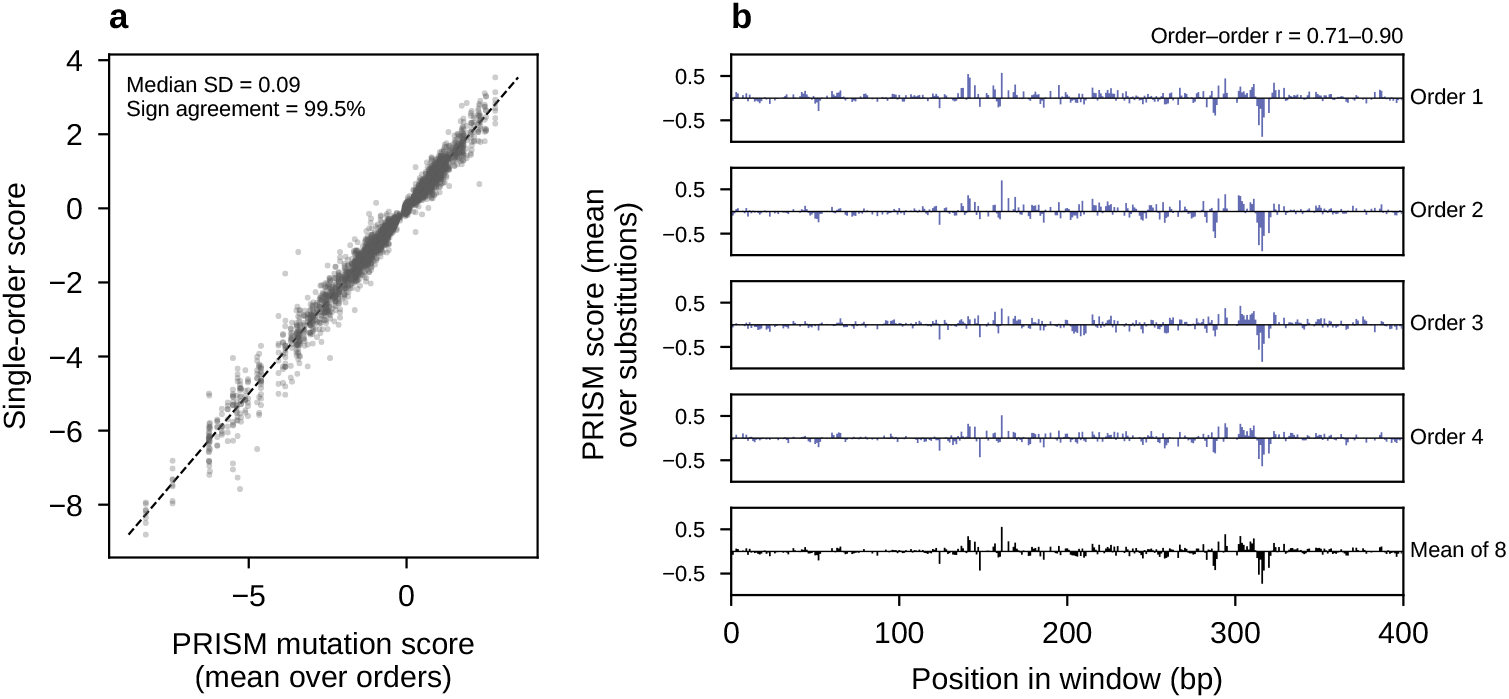
Sensitivity of PRISM scores to the unmasking order. **a** Scores of each mutation selected in the PRISM validation (observed target reads; Supplementary Note 6.1) under each of eight independent confirmation orders, against their mean over the eight orders. The dashed line is *y* = *x*. Median SD is the median over mutations of the standard deviation across orders, and sign agreement is the fraction of single-order scores with the same sign as the mean, for mutations whose mean exceeds 0.05 in magnitude. **b** PRISM scores in the 400-bp window of one randomly chosen observed target read, averaged over alternative bases at each position, for four of the eight unmasking orders and the mean over all eight. Positions without substitutions that preserve the SMF position mask were not scored and are left blank. Order–order *r* is the range of pairwise Pearson correlations between orders across all substitutions.

The empirical effect of a mutation on a target read *x* is

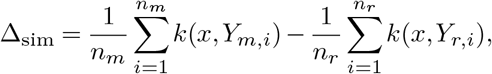

where *k*(*x, Y*) is the fraction of the target’s valid positions at which a generated read *Y* has the same methylation call, and *Y*_*m,i*_ and *Y*_*r,i*_ are the *n*_*m*_ = *n*_*r*_ = 1,024 reads generated from the mutant and reference sequences.

### Supplementary Note 7: Interpretation of the compositional accessibility baseline

The Methods define a compositional accessibility baseline for each combined motif construct BAB, in which a central motif, *A*, sits between two copies of a partner motif, *B*. We write *F* for the two flanking copies of *B* taken together. The baseline is built from the accessible fractions of the empty background (− − −), the central motif alone (−A−), and the paired flanks alone (B−B), which we denote *p*_0_, *p*_*A*_, and *p*_*F*_, so that the subscript names the motifs present relative to the background. Here we explain the reasoning behind its construction and how departures from it should be interpreted. In the notation of the Methods, for *X ∈ {A, F }*, the baseline is

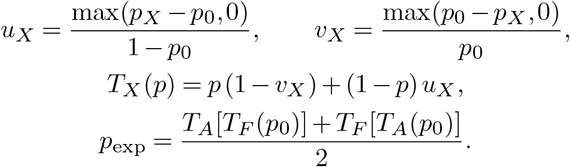

Accessibility is defined using the same central window and molecule-filtering criteria in all four constructs. Because *F* includes both flanking motifs, its response can already include interactions between them.

#### Construction

An accessibility gain is normalized by the background inaccessible fraction, 1 *−p*_0_, whereas a loss is normalized by the background accessible fraction, *p*_0_. For example, an increase from 20% to 60% accessibility corresponds to a gain of 40 percentage points, or half of the background inaccessible fraction, *u*_*X*_ = 0.40*/*0.80 = 0.50. *u*_*X*_ captures a net gain and *v*_*X*_ a net loss in accessibility, so at most one of them is nonzero. In *T*_*X*_ (*p*), the first term retains the accessible fraction after applying the normalized loss, and the second adds the normalized gain from the inaccessible fraction. By construction, *T*_*X*_ (*p*_0_) = *p*_*X*_, so applying the transformation to the background reproduces *p*_*X*_ . The output of *T*_*X*_ also stays between zero and one.

Averaging over both orders makes the expectation symmetric with respect to the central motif and the paired flanks. The two orders give the same result when *A* and *F* both increase accessibility or both decrease it, but can differ when one increases accessibility and the other decreases it. This averaging is a mathematical convention for combining their effects and does not specify the biological order of TF binding or chromatin-state transitions. Likewise, *u*_*X*_ and *v*_*X*_ summarize accessibility differences between constructs and are not measured transition rates.

**Supplementary Table 2.**
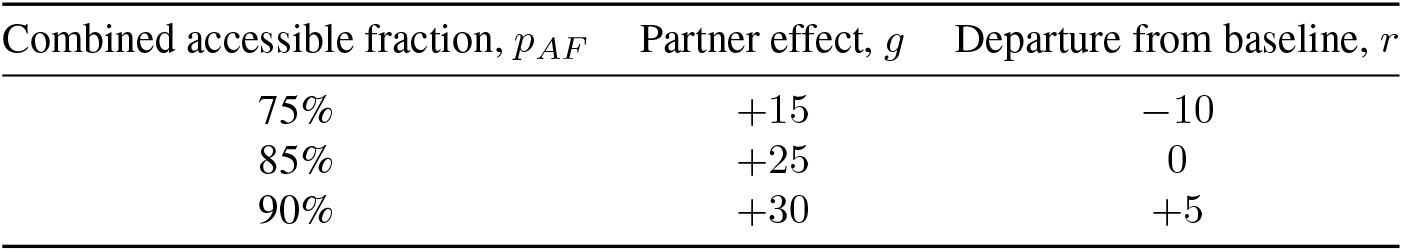
Illustrative outcomes for the example with *p*_0_ = 0.20, *p*_*A*_ = 0.60, and *p*_*F*_ = 0.70, for which *p*_exp_ = 85%. Differences are in percentage points.

#### Overlap between gains

When *A* and *F* both increase accessibility, writing their gains as Δ_*A*_ = *p*_*A*_*− p*_0_ and Δ_*F*_ = *p*_*F*_ *−p*_0_ gives

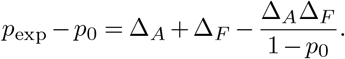

The final term accounts for overlap between the accessibility gains attributed to *A* and *F* . Thus, the expected combined gain is smaller than Δ_*A*_ +Δ_*F*_ whenever both are positive. This correction applies even when *p*_0_ +Δ_*A*_ +Δ_*F*_ is below one, so it is not simply a cap at 100% accessibility.

An illustrative example makes the overlap assumption explicit. Suppose that *p*_0_ = 0.20, *p*_*A*_ = 0.60, and *p*_*F*_ = 0.70. The baseline gives

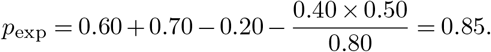

In a hypothetical population of 1,000 molecules, 200 would be accessible in the background. The central motif and the paired flanks would each make some of the remaining 800 molecules accessible, 400 and 500 molecules, respectively, which we call their response subsets. The baseline corresponds to an expected overlap between the two response subsets of

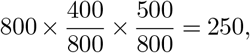

giving 200 + 400 + 500 *−*250 = 850 accessible molecules. This construction assumes that the two response subsets overlap according to their frequencies within the background-inaccessible population. It does not assume independent TF binding. These subsets are only an illustration, because reads generated for different constructs are not the same physical molecules.

#### Partner effects and baseline departures

We distinguish the partner effect, the effect of adding the paired flanks to the central motif, from the departure from the baseline. Within a matched background, these quantities are

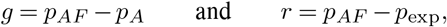

respectively, where *p*_*AF*_ is the accessible fraction of the combined construct. The partner effect *g* measures whether adding the paired flanks increases or decreases the accessible fraction relative to the central motif alone. The baseline departure *r* measures whether the combined accessible fraction is greater or smaller than expected from the composition rule. These comparisons answer different questions and can have different signs. With the same *p*_0_, *p*_*A*_, and *p*_*F*_ as in the example above, the combined construct could take any of the values in Supplementary Table 2.

A combined accessible fraction of 75% therefore represents an increase relative to the central motif alone, despite falling below the 85% baseline. In terms of the response subsets, this outcome could arise if the two subsets overlapped on 350 molecules rather than the expected 250. Conversely, 90% combined accessibility could arise with an overlap of 200 molecules. These examples show that departures of either sign can arise from different degrees of shared responsiveness, without requiring inhibition or enhancement of TF binding.

#### Interpreting departures

A negative departure indicates less accessibility than the composition rule predicts, a positive departure indicates more accessibility, and a departure near zero indicates agreement within estimation uncertainty. These outcomes do not, by themselves, establish antagonistic, cooperative, or independent TF binding. The responses to *A* and to *F* can already reflect TF–nucleosome interactions, and the paired-flank response can include cooperation between its two sites. Agreement with the baseline therefore does not imply an absence of cooperativity. Departures identify limitations of the composition rule, and their causes could include shared sequence dependencies, changes in TF–nucleosome competition or nucleosome positioning, direct TF interactions, or prediction and state-calling errors. Distinguishing these explanations requires additional occupancy measurements, perturbations, or explicit mechanistic models. The bootstrap confidence intervals for the baseline comparisons (Fig. 4b and Extended Data Fig. E7a–b) describe uncertainty from the sampled backgrounds conditional on the fitted SMFNet model, and they do not quantify uncertainty in the model itself or provide independent biological validation.

